# What and where manifolds emerge and align with perception in deep neural network models of sound localization

**DOI:** 10.64898/2026.02.11.705311

**Authors:** Chenggang Chen, Zhiyu Yang, Xiaoqin Wang

**Affiliations:** Department of Biomedical Engineering, Johns Hopkins University

## Abstract

Whether the auditory cortex has parallel pathways for sound identification (“what”) and localization (“where”), and whether it contains a map of auditory space, is debated. Here, we examined the low-dimensional structure (manifold) of “what” and “where” representations in deep neural network models of sound localization. Unexpectedly, models trained for “where” learned untangled “what” manifolds, including voice type, reverberation, and spectral detail. The distribution of “what” manifolds was not random, but geometrically organized by spectral similarity. The separability and distance of both “what” and “where” manifolds were aligned with human behavior. “What” also determined whether “where” manifolds organized into a map: maps emerged when “what” contained localization cues that were topographically organized. However, forming a spatial map reduced localization accuracy in both models and human listeners. Together, object manifolds reveal learned task-irrelevant attributes of object that are ignored when measuring task performance alone, and task-optimized neural networks can provide insights into brain and behavior, not just replicate them.

## 1 Introduction

A central goal of neuroscience is to build models that reach human-level performance on perceptual tasks, reproduce brain responses, and, more importantly, explain the neural mechanisms. The hierarchical nature of sensory systems has established deep neural networks (DNNs) as the leading models in the field which transform sensory inputs into internal representations (Yamins et al., 2014; Kell et al., 2018; Keshishian et al., 2020; Millet et al., 2022; Vaidya et al., 2022; Giordano et al., 2023; Drakopoulos et al., 2025). Deep neural network (DNN) models of sensory systems have mainly focused on task performance and neural activations, largely neglecting object manifolds—low-dimensional structures embedded within the high-dimensional space of neural activations, formed by population responses to the same object under varying conditions (e.g., a cat image at different angles) (Chung & Abbott, 2021). The original dimensionality corresponds to the number of simultaneously recorded neurons in the brain and the number of convolutional kernels or units in DNNs. In the brain, neural manifolds have been widely used to characterize cognition (Chaudhuri et al., 2019; Langdon et al., 2023), object recognition (DiCarlo & Cox, 2007), neural dynamics during movement (Perich et al., 2025), and to achieve long-term (Chen et al., 2024) and cross-subject (Safaie et al., 2023) movement decoding. In DNNs, object manifolds are used to check representations of image (Kazemian et al., 2025), audio (Chen & Yang, 2025), and in unsupervised/self-supervised learning (Zhuang et al., 2021). Although the geometry of the object manifold in DNNs depends on layer (Cohen et al., 2020) and category (Doshi & Konkle, 2023; Zhang et al., 2024b), it remains unknown whether object manifolds in DNN models of sensory systems are task-specific and behavior-aligned.

Unlike hypothesis-driven and hand-crafted models with task-specified parameters, data-driven and hypothesis-free DNN models of sensory systems, like our brains, take real-world sensory inputs. For instance, a face carries both viewpoint and identity attributes, just as a voice contains both pitch and location. In earlier cortical face-processing pathway, same neurons are tuned to both face viewpoint and identity (Freiwald & Tsao, 2010). When DNN is trained on a single task-relevant attribute, task-irrelevant attributes remain inseparable in the inputs. Consequently, it remains to be tested whether manifolds for these task-irrelevant attributes emerge within DNN models of sensory systems.

Here, we used auditory parallel pathways as a testbed to characterize the object manifolds of task-relevant and task-irrelevant sensory attributes in DNN models of sensory systems. In the primate visual cortex, information is processed in a hierarchical manner using two parallel pathways (Mishkin et al., 1983): the ventral, or “what” pathway, and the dorsal, or “where” pathway (Fig. 1a). These two pathways are specialized for visual identification/categorization and localization/movement, respectively. The existence of parallel pathways in the auditory cortex (AC) is highly debated, when it was first proposed around the 2000s (Rauschecker & Tian, 2000) (Fig. 1a). Anatomical studies found that caudal and rostral streams of auditory afferents target dorsal and ventral domains in the macaque monkey prefrontal cortex (Romanski et al., 1999). Neurophysiology study in anesthetized macaque also found that single neurons in the caudal AC were highly tuned for sound locations, whereas neurons in the rostral AC are more selective for conspecific vocalization (Tian et al., 2001). Functional magnetic resonance imaging (fMRI) in humans also revealed a similar dichotomy in AC. Behavior studies in cats and humans further show causality of modulation of these two streams in what and where discrimination tasks (Lomber & Malhotra, 2008; Ahveninen et al., 2013). On the other hand, the theory of auditory parallel stream has been argued since it was first proposed (Belin & Zatorre, 2000; Hall, 2003). Multiple pieces of evidence are against the parallel stream hypothesis. First, the distribution of “what” is everywhere since neurons in both caudal and rostral areas of awake macaque auditory cortex are equally selective for vocalization (Recanzone, 2008; Bizley & Walker, 2009). Second, although neurons in the caudal AC are more selective for sound locations, highly spatially tuned neurons were also identified in the rostral AC (Woods et al., 2006; Remington & Wang, 2019). Third, AC neurons show multiplexed representation of sound features, i.e., “where” (location) and “what” (pitch and timbre) information (Bizley et al., 2009; Walker et al., 2011). Last, behavior studies using the advanced optogenetics tool show that inhibiting one area of AC impaired both spatial and non-spatial hearing (Town et al., 2023).

**Figure 1:**
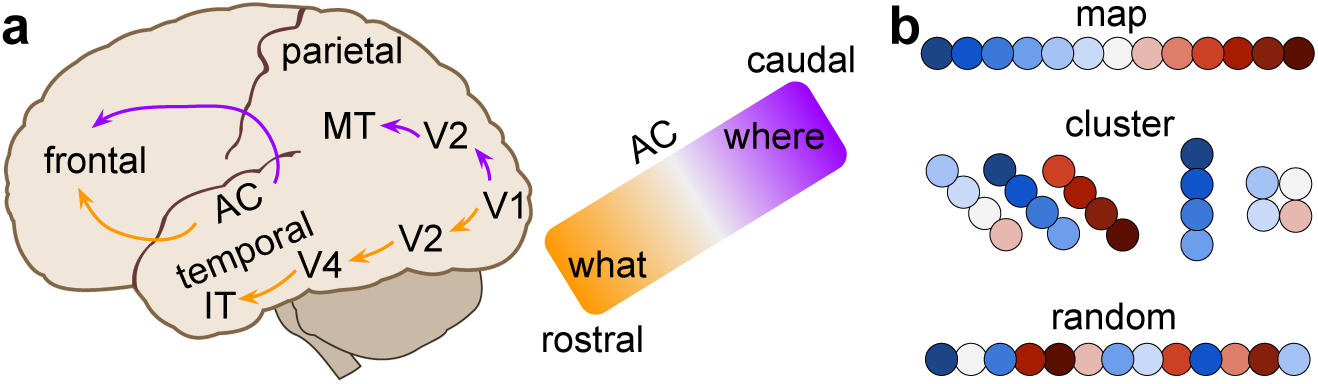
**a**. In visual cortices, the dorsal “where” pathway (purple arrows) starts from the primary visual cortex (V1), and projects to V2 and middle temporal (MT) cortex in the parietal lobe. The ventral “what” pathway (orange arrows) also starts from V1, then projects to V2, V4, and inferior temporal (IT) cortex in the temporal lobe. In the auditory cortex (AC), caudal “where” and rostral “what” pathways project to dorsal and ventral frontal cortex, respectively. **b**. Representations of sound locations (color dots) formed three candidate organizations in the brain. Top: topographic space map. Middle: neighboring neurons on the same electrode from tangential penetration, normal penetration, and from different electrodes but closely spaced normal penetration show similar spatial tuning. Bottom: two-photon calcium imaging (Panniello et al., 2018) and normal electrode penetration (Remington & Wang, 2019) found neighboring neurons have diverse spatial tunings.

What makes the above arguments of auditory parallel streams more complex is the other mystery about the auditory space map in AC (Fig. 1b). According to the parallel pathway, spatially tuned neurons should form an orderly space map in the dorsal AC. Neurons selective for sound types should form several clusters in the rostral AC, like the ones in the visual ventral pathway (Bao et al., 2020). Although there is some evidence of voice patch existence in rostral AC (Petkov et al., 2008), no evidence exists for a space map in any area of AC (Middlebrooks, 2021). A neural map of auditory space was first found in the barn owl (Knudsen & Konishi, 1978), then in the mammalian inferior/superior colliculus (IC/SC) (Chen & Song, 2024; Bianchini et al., 2025). Middlebrooks and colleagues and other groups started to search for such a map in AC since 1981 in cats (Middlebrooks & Pettigrew, 1981). After 40 years of research, there is still no evidence for such a map in the AC for all the species that have been examined (Middlebrooks, 2021). Unlike the visual and somatosensory systems with orderly mapped receptors in the retina and skin, the cochlea has a map of sound frequency instead of location. Since the sound locations were computed inside the central nervous system, the representations of those computed but not relayed locations could be in any structure. After two decades of research on the parallel pathway (Rauschecker & Scott, 2009; Recanzone & Cohen, 2010), and four decades of research on the space map, our understanding of these two questions is still very limited, partially due to that we lack a computational model for them.

There are many models about either sound spatial and nonspatial attributes. For example, the most famous delay line model (Jeffress, 1948) for computing interaural time difference (ITD), and the spectral-temporal receptive field (STRF) model (Theunissen et al., 2000) for characterizing the content of natural sound. Although mechanistic models are useful for explaining how individual neurons are tuned to spatial and nonspatial sound features, they cannot model a spatial organization like parallel streams or a space map. The self-organizing map (SOM) has been used successfully to model orientation and direction maps in the visual cortex (Durbin & Mitchison, 1990). Because their inputs are only hand-crafted simple features like line directions, SOM could not take inputs from two different sound attributes or compute the ITDs. We have made three contributions in this study.

We built the first computational models for both parallel pathways and space maps in the auditory system. Our audio datasets contain both “what” (content) and “where” (location) attributes of sounds. Our models are hypothesis-free, task-optimized DNNs trained solely for sound localization. Our model inputs include both binaural audio waveforms of animal vocalizations and cochleagrams of speech filtered by the pinnae, head, and torso from real-world human listening environments.

We are the first to examine the separability and geometry of object manifolds from DNN models of the human auditory cortex or human listening task. Previous studies either compare the activations of model units and human auditory cortex to the same sound stimuli (Kell et al., 2018; Li et al., 2023; Tuckute et al., 2023), or compare the task performance between models and humans (Saddler et al., 2021; Francl & McDermott, 2022; Feather et al., 2023; Saddler & McDermott, 2024).

We are also the first to study the neural mechanisms underlying DNN models of human sound localization behavior. Three previous studies (Francl & McDermott, 2022; Saddler & McDermott, 2024; Banerjee et al., 2025) showed that their models exhibited many features of human spatial hearing based on task performance. By examining the task-relevant and task-irrelevant manifolds, our work fills the gap between DNN models of human auditory perception and auditory systems.

## 2 Results

Here, we modeled the representation of sound locations with a DNN trained for the sound localization task. We used an open-source bioacoustics sound localization dataset (Peterson et al., 2024). Sounds were played from 394 locations at the bottom of an arena (Fig. 2a) and captured by four microphones at the corners that were 35 cm from the bottom. Each location has a median of 144 stimuli, including a fixed number of six sound types and three sound levels, and eight (median) different samples of the same sound type. Sound types include five different classes of gerbil vocalization and one artificial white noise (Fig. 2b). Notice that “dfm”, “sc”, and “stack” calls have equally spaced frequency bands in their spectrograms. The first or bottom frequency band has a fundamental frequency of f0. Other bands are called harmonics and have a frequency that is an integer multiple of f0. For example, there are three, two, and five harmonics in these three sound types. “upfm” and “warble” only have one narrow band of frequency modulations. In contrast, sound energy was distributed uniformly in the “white” noise call. Notice that unlike spectrograms, the audio waveforms are very different from each other (Supplementary Fig. 1a-c). The medium sound level is calibrated to be approximately the same level as a natural vocalization. The soft and loud sound levels are around 6 dB lower or higher than the medium level (Fig. 2c). We fed pairs of raw audio waveforms into a deep neural network (DNN) consisting of five one-dimensional convolutional layers (Fig. 2d), trained exclusively on a sound localization task. From the trained model, we extracted high-dimensional representations from each layer (Fig. 2e) to analyze their geometric properties.

**Figure 2:**
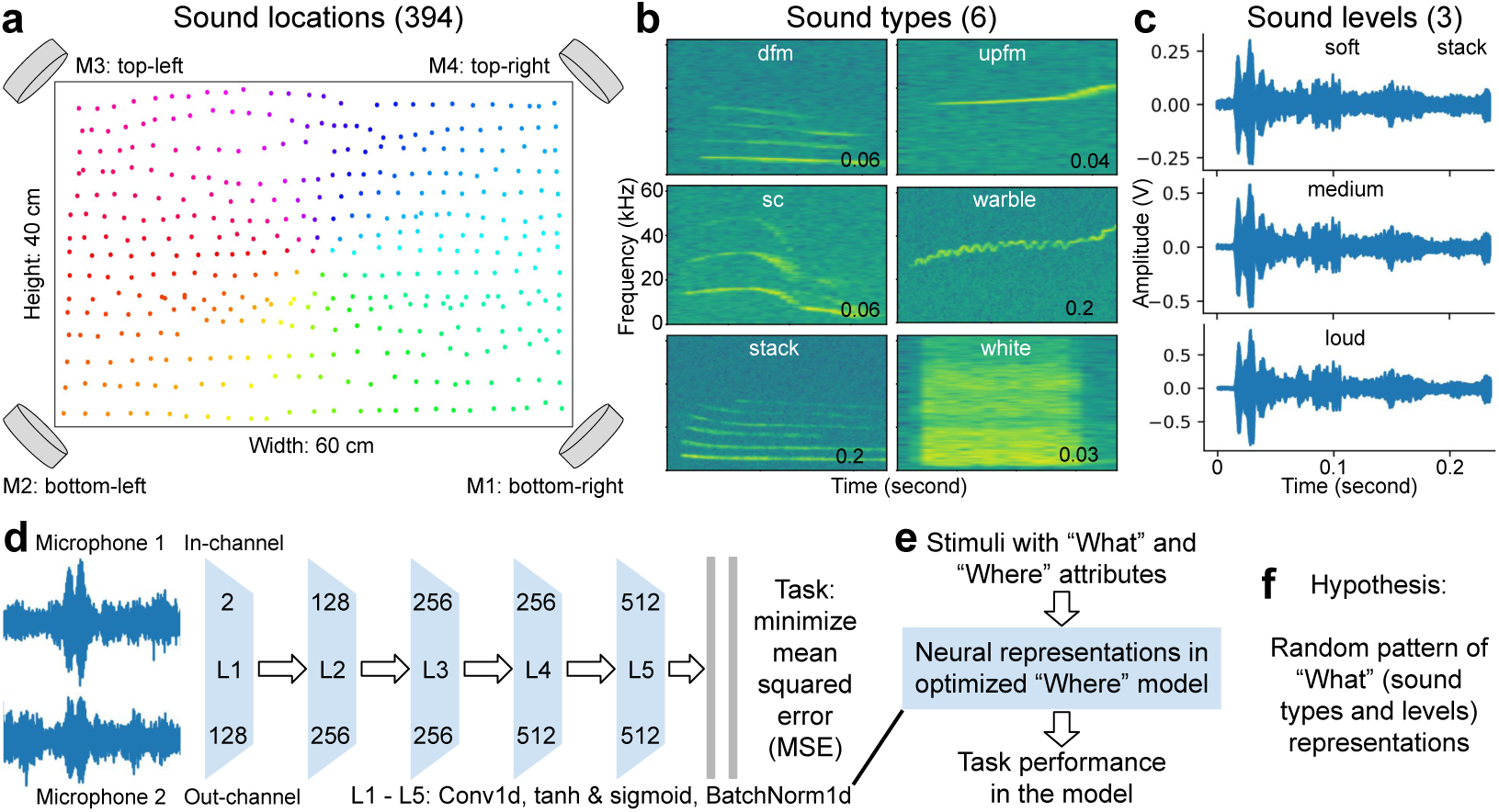
**a**. Four microphones at the corners of the arena will record sounds coming from 394 locations. **b**. Spectrograms of six example sound types. Here, “fm” stands for frequency modulation, “d” for downwards, and “sc” for soft chirp. **c**. Three different sound levels for the call type “stack”. **d**. Model architectures. **e**. Pipeline to extract the neural representations. **f**. Our hypothesis.

To quantify the dimensionality of these representations, we employed two complementary metrics (see Methods). First, we calculated the number of principal components required to explain a sufficient threshold (e.g., 90%) of the variance (Cohen et al., 2020). This method identifies dimensionality via the “elbow point” in the variance histogram, similar to principal component analysis (PCA). Second, we calculated the effective dimensionality, also known as the participation ratio (PR) (Gao et al., 2017), defined as ( *λ_i_*)^2^*/* (*λ*^2^), where *λ_i_* represents the eigenvalues of the covariance matrix of the neural population activity. Following a recent critique by Chun et al. (2025) that the standard PR is biased downwards by small sample sizes, we utilized their proposed bias-corrected estimator to ensure accuracy under finite sampling conditions. We measured the dimensionality of sound type and sound level manifolds individually (Supplementary Fig. 1d). Analysis of the explained variance (*R*^2^) revealed “elbow” drops at the 4th and 3rd components for sound type and level, respectively, accounting for 57% and 48% of the cumulative variance. However, reaching a 90% variance threshold required significantly higher dimensions (sound type: 36, level: 60) due to the slow decay of the variance curve after the first ten components. In contrast, the PR yielded much lower effective dimensionalities (sound type: 9.4, level: 14.6). Notably, the bias-corrected PR produced nearly identical results (e.g., 9.39 for sound type), confirming the robustness of these estimates. These results demonstrate that in the trained DNN, individual stimulus representations reside on manifolds of significantly lower dimensionality than the ambient space (512D).

To visualize these low-dimensional manifolds, we employed uniform manifold approximation and projection (UMAP) (McInnes et al., 2018). We selected a 2D projection for clarity, comparability to cortical surface maps, and its ability to preserve over 40% of the variance. As demonstrated in subsequent sections, our quantitative findings remained highly consistent between the projected 2D space and the full 512D representations (Figs. 5c and 6e). Given that the model was trained solely to localize sound sources (“where”), we hypothesized that its representations of sound type and level (“what”) would not be explicitly untangled, resulting in random patterns (Fig. 2f).

Surprisingly, despite being trained exclusively for sound localization, the DNN spontaneously organized representations according to sound type (Fig. 3a). We observed that sound types sharing similar spectral characteristics formed distinct clusters, suggesting the model learned to group inputs by spectrogram similarity even though it received raw waveforms. Specifically, “warble” and “upfm”, two sound types characterized by a single narrow frequency band, clustered in close proximity. Their corresponding localization representations reside on low-dimensional manifolds (PR: 7 and 11, respectively) but appear overlapped in the 2D projection (Fig. 3c). Similarly, the three harmonic sound types (“dfm”, “stack”, and “sc”) formed a cohesive group and also occupied lowdimensional manifolds (PR: 11, 10, and 9), though they generally overlapped in the 2D space. The white noise cluster (PR: 8), characterized by broad frequency bands, was situated adjacent to the harmonic group but distinct from the narrow-band types. Notably, the white noise representation formed a clear, organized spatial map (Fig. 3c). In contrast, representations of sound levels (loud, medium, soft) were globally entangled, largely following the geometry of the sound type clusters (Fig. 3b). However, their local geometry remained preserved within a relatively low-dimensional subspace (PR: 14, 15, and 15). When comparing the two features, the variability of sound level representations was significantly smaller than that of sound types (0.16 vs. 2.14). Thus, manifolds for both sound type and level emerge spontaneously in this “where” model, with sound type manifolds exhibiting lower dimensionality but higher variability compared to those of sound levels.

**Figure 3:**
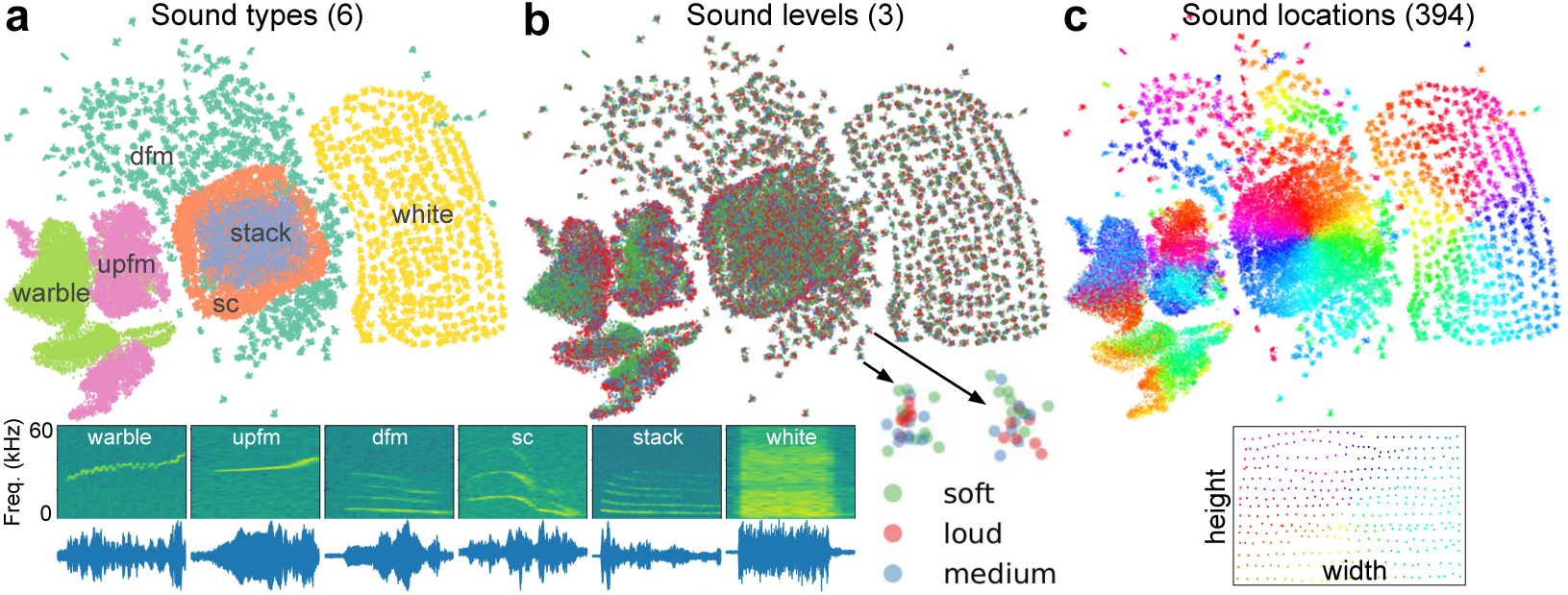
**a-c**. Object manifolds of sound types, sound levels and sound locations. Supplementary Fig. 2 and 4a show the manifolds across all five and ten layers, respectively.

To quantify the differences in clustering between layers, we used the normalized mutual information (NMI). NMI is an information-theoretic metric used to evaluate the similarity between two clusters. The NMI ranges from 0 to 1, with 0 representing no mutual information. Supplementary Fig. 2a-c show the representations of all three sound attributes (rows) across all five layers (columns) in two microphones. Since the representations of sound levels are random globally, their NMIs were near 0. The sound type representations were not well separated in layer 1 (NMI: 0.54), as two sound types overlapped and all clusters were close to each other. The NMIs increased from layer 2 (0.61) to peak at layer 3 (0.89) and then decreased to 0.62 in layer 5. The “white” noise type of sound began to show a twisted map of sound location from layer 2. After layer 3, the representations became scrambled again to better classify sound locations. The map for “white” noise was clear with high separability. In contrast, the maps for “sc” and “stack” has low separability. We observed similar patterns in the other two microphones (Supplementary Fig. 2d-f). We also computed NMIs in the original 512D space without using UMAP and found very similar results.

To quantify the organization strength of the neural representations, we calculated the explained variance (*R*^2^) from a linear regression between the 2D UMAP coordinates and the ground-truth 2D sound locations (azimuth and elevation). We observed striking differences in organization strength between sessions; for example, one session exhibited a clear topographic map with high organization (*R*^2^ = 0.99), while another lacked clear structure (*R*^2^ = 0.22) (Supplementary Fig. 3a, b). Surprisingly, despite this structural disparity, the sound localization performance was comparable between the two sessions (MSE: 0.31 vs. 0.37 cm). Extending this analysis across all six microphone pairs and five stimulus types (Supplementary Fig. 3c-d), we found no significant correlation between organization strength and task performance (*p* = 0.202*, R*^2^ = 0.14).Importantly, our model is entirely task-driven and hypothesis-free: it was trained strictly to localize sound, with no constraints imposed to enforce clustering or topographic map formation.

As a control, we repeated the regression analysis using the full 512D representational space against the 2D sound locations (Supplementary Fig. 3e). In this high-dimensional space, the correlation with task performance was significant (*p* = 0.001*, R*^2^ = 0.39), driven by the fact that all sound types exhibited high representational fidelity (*R*^2^ *>* 0.89). This control confirms that the weak organization observed in the 2D projection for some conditions was not an artifact of insufficient training (since the 512D representations were highly accurate) nor strictly a failure of the dimensionality reduction (since over half of the sound types maintained high *R*^2^ in both spaces). We further examined whether the intrinsic dimensionality of the manifolds predicted performance. We compared task performance against both the dimension explaining 90% of variance and the bias-corrected participation ratio (Supplementary Fig. 3f). Neither metric showed a significant correlation with performance (*p >* 0.1; *R*^2^ = 0.15 and 0.04, respectively). These results suggest that while all sound type representations reside on low-dimensional manifolds (relative to the 512D ambient space), the specific dimensionality of these manifolds is irrelevant to localization accuracy. This reinforces our finding that topographic maps, such as those visible in 2D UMAP, are not a prerequisite for high performance. In summary, a DNN trained solely on sound localization spontaneously generates a diversity of representational geometries, ranging from topographic maps and clusters to seemingly random patterns, mirroring the diversity observed in the SC/IC and AC. These findings demonstrate that a topographic map is not the unique or optimal solution for accurate sound localization.

We observed consistent findings when using a deeper network with ten (instead of five) layers (Supplementary Fig. 4a). Two sound types (“upfm” and “warble”) that are partially overlapping in layer 1 keep this organization all the way to layer 10. In contrast, the other four sound types are only roughly separated in layer 1 and become gradually well separated in deeper layers. The “white” noise type of sound begins to show a spatial map after layer 3 and reaches a peak at layers 6–7. The organization is further refined, with organized clusters formed at each location. The “sc” sound type begins to show maps at layer 7 and reaches its peak at layer 9. We quantified the clustering accuracy (NMI) and organization strength (*R*^2^) across all ten layers using different values of UMAP’s main hyperparameter, the number of neighbors. Except for the two smallest neighbor values at very deep layers, the NMI values are reliable across choices of neighbors (Supplementary Fig. 4b). The *R*^2^ scores are always highest for the “white” noise and “sc” sound types and peak in the middle to deep layers, regardless of the number of neighbors (Supplementary Fig. 4c). Therefore, our findings hold for both shallow and deeper network models and are robust to changes in UMAP hyperparameters.

**Figure 4:**
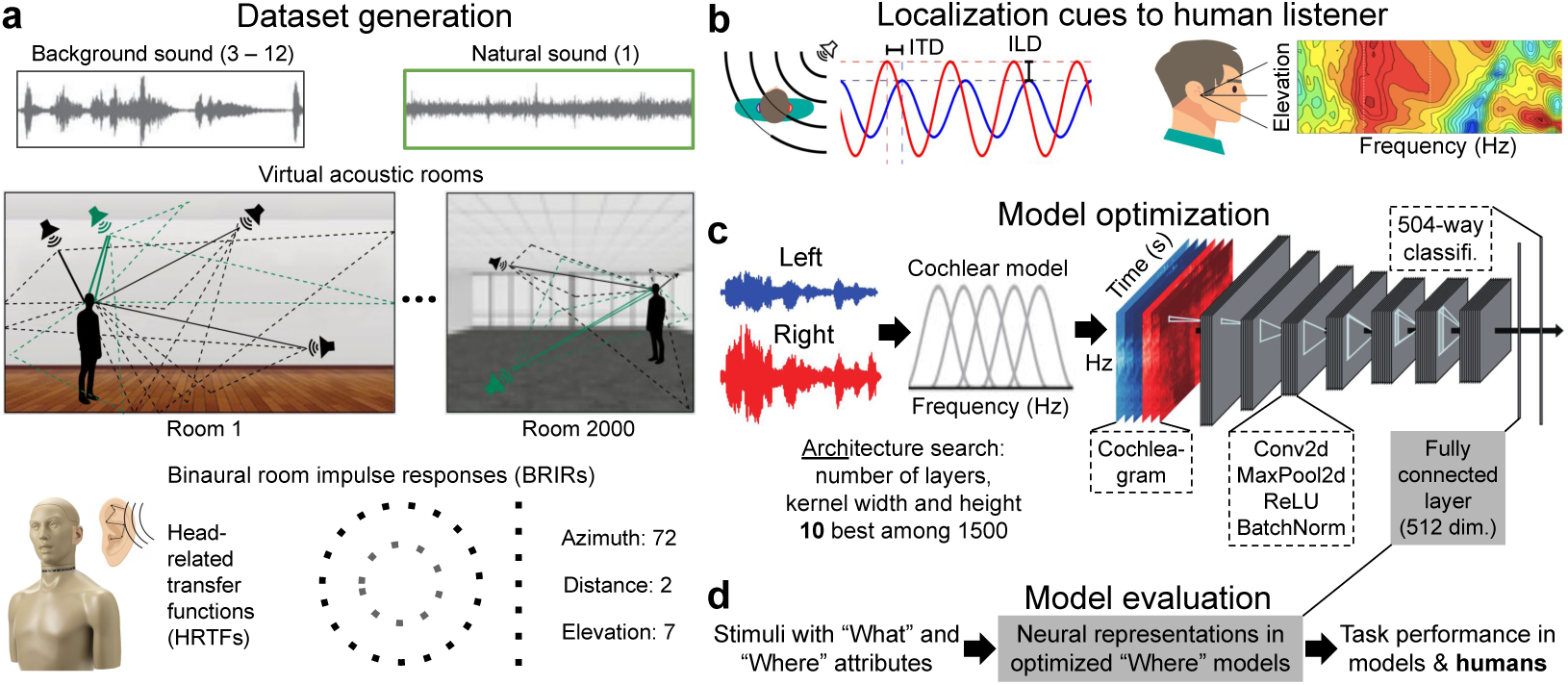
Sound localization dataset and models for human listeners. **a**. Natural sound (green) is rendered at one location and multiple background sounds (black) are rendered at other locations. There are 2000 different simulated rooms with different sizes and floor and ceiling materials (copied from Fig. 1c of Francl & McDermott (2022)). In addition to the rooms, rendering also includes direction-specific filtering by the head/torso/pinnae, using the HRTFs from the KEMAR manikin (copied from the website of G.R.A.S Acoustics). **b**. Sound localization cues available to human listeners. Left: inter-aural time and level differences (ITDs and ILDs) (copied from Fig. 3b of Saddler & McDermott (2024)). Right: monaural spectral cues to sound elevation. Color-coded HRTFs (amplitude spectra (1–16 kHz) between -15 dB (dark blue) and 20 dB (dark red)) are shown as a function of elevation for a typical human subject (copied from Zonooz et al. (2019)). **c**. Localization model schematic (modified from Fig. 3a of Saddler & McDermott (2024)). **d**. Sound stimuli are fed into optimized “where” models, from which 512D activations are extracted from the penultimate layer. The geometric properties of representations are then compared against task performance.

**Figure 5:**
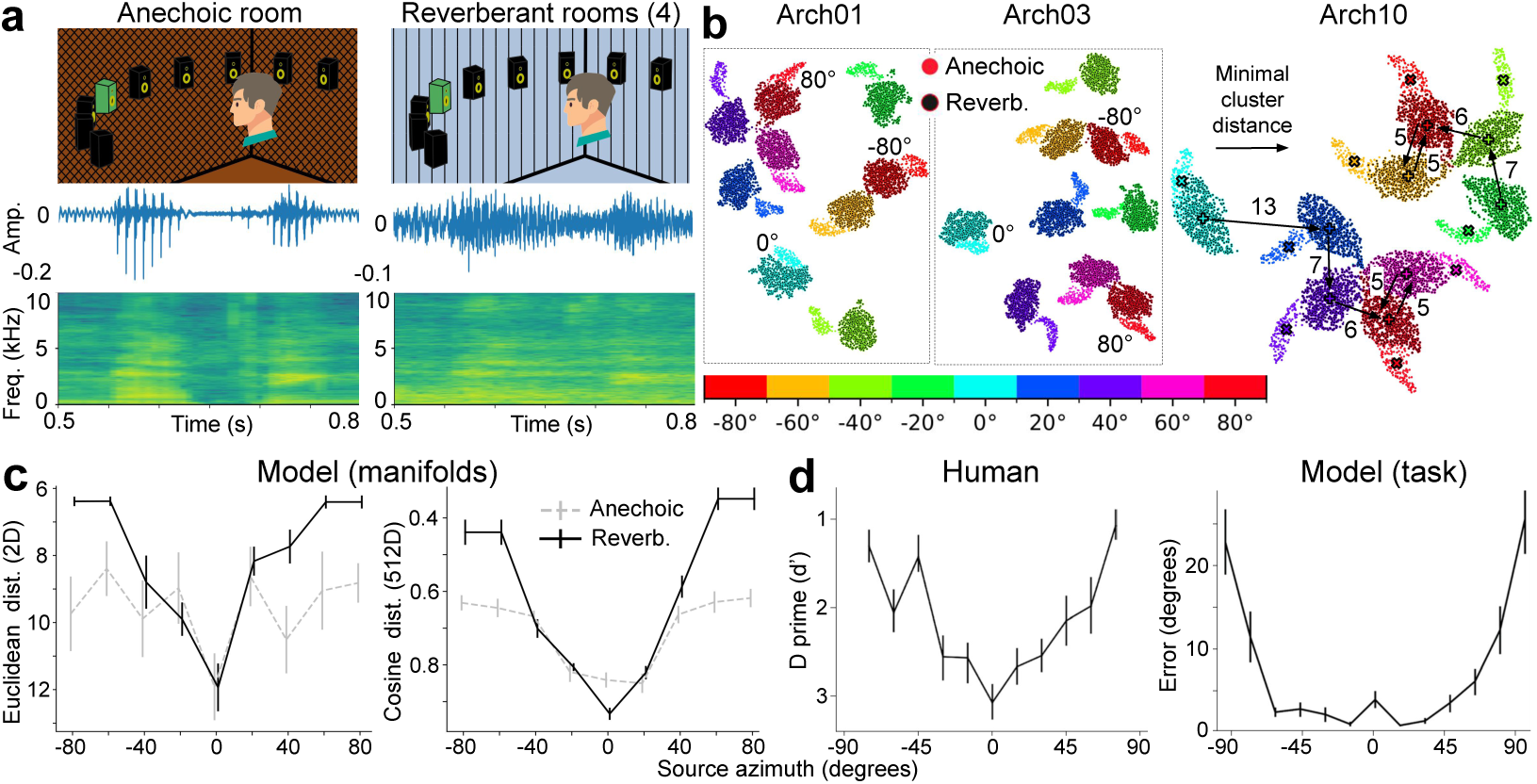
Object manifolds are sensitive to reverberation and display the highest separation at the midline, similar to human listeners. **a**. Top: schematic of the sound localization experiment in two conditions. There are four types of reverberant rooms (copied from Fig. 7d in Saddler & McDermott (2024)). Bottom: sound waveforms and spectrograms of the same speech segment. **b**. Manifolds of nine azimuths in the two room conditions (anechoic: colored dots; reverberant: colored dots with black centers). Right: minimal cluster distances (values next to arrows) of manifolds at seven azimuths in reverberant rooms for Arch10. Black “+” or “x” indicates manifold centers. **c**. Minimal cluster distances in 2D space (Euclidean, left) and native 512D space (Cosine, right). Error bars: SEM (n=10 models). Number of neighbors of UMAP=50. **d**. Localization accuracy of the human listeners (left) and models (right) for broadband noise at different azimuths (copied from Fig. 3b, c in Francl & McDermott (2022)). Error bars: SEM (n=12 human listeners and 10 models).

**Figure 6:**
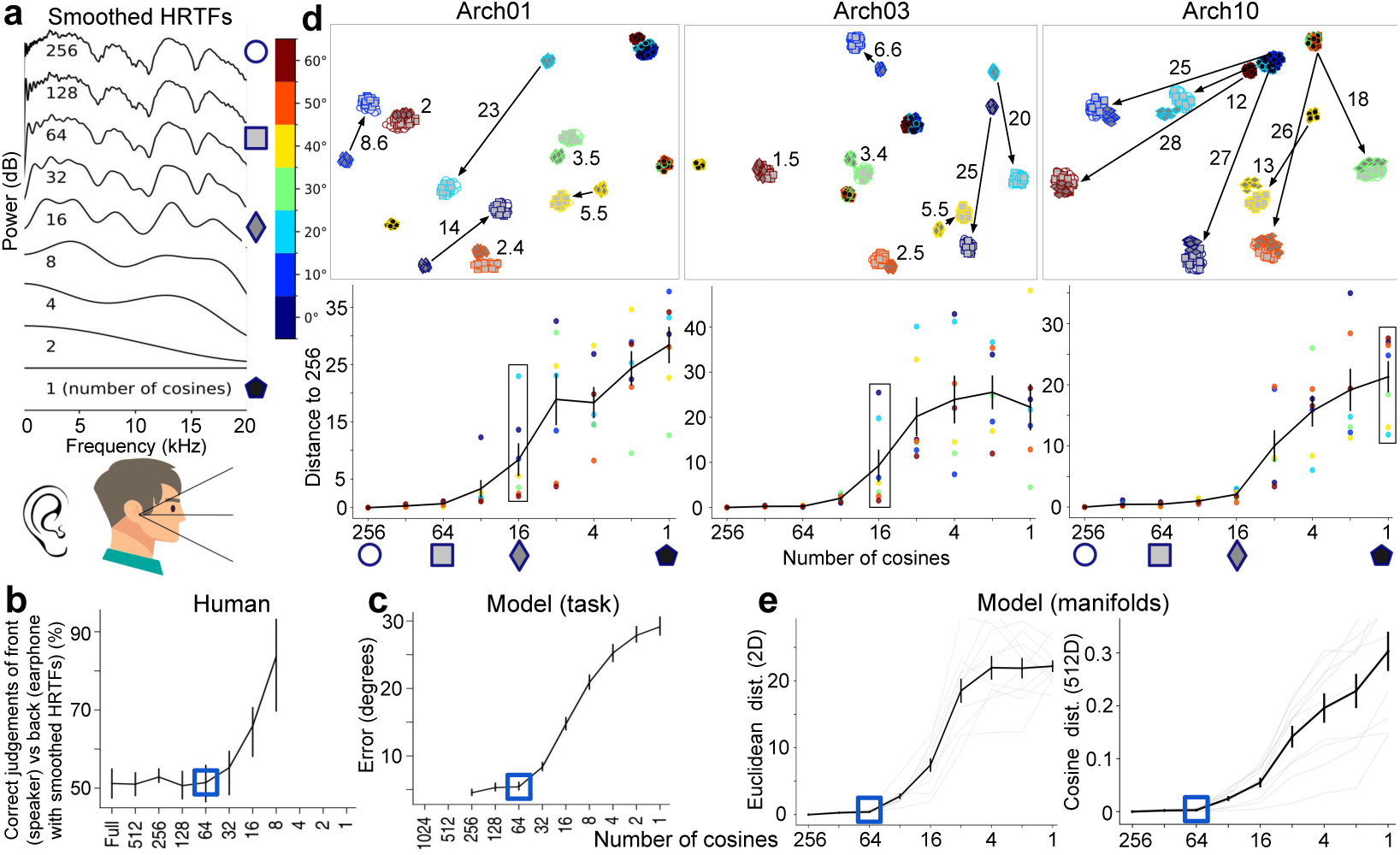
Object manifolds of sound elevations exhibited similar spectral resolution to human listeners. **a**. Smoothing the power spectra of HRTFs by lowering the number of cosines used to approximate the discrete cosine transform (copied from Supplementary Fig. 3d in Saddler & McDermott (2024)). **b**. Effect of spectral smoothing on human perception. Participants heard two sounds, one played from a speaker in front of them and one played through open-backed earphones, and judged which was which. The earphone-presented sound was rendered using HRTFs smoothed to varying degrees. In practice, participants performed the task by noting changes in apparent sound location. The figure is copied from Fig. 4i in Francl & McDermott (2022). **c**. Effect of spectral smoothing on model’s sound localization accuracy (measured in both azimuth and elevation ). The figure is copied from Fig. 4j in Francl & McDermott (2022). **d**. Top: Elevation manifolds visualized across four spectral densities. Arrows indicate the distance between the depicted condition (1 or 16 cosines) and the 256-cosine baseline. Bottom: Distance to the 256-cosine baseline as a function of spectral density for seven elevations. Black squares correspond to the conditions shown above. Error bars: SEM (n=7 elevations). **e**. Distance to the 256-cosine baseline quantified in the 2D space (Euclidean, left) and native 512D space (cosine, right). Error bars: SEM (n=10 models).

Our current models take stereo audio waveforms but does not consider the acoustic cues created by direction-specific filtering of the animal/human body, nor the cochlear properties of the animal/human. Therefore, the models we use here bear little resemblance to the architectures of biological, especially human, auditory systems. It is also unclear whether our findings (low-dimensional “what” representations or manifolds emerge in the “where” model and that a space map is unnecessary) hold for datasets other than gerbil vocalizations. To address the limitations in our models and datasets, we analyzed the neural representations of sound locations in models optimized for human sound localization behaviors (Francl & McDermott, 2022; Saddler & McDermott, 2024). These task-optimized models (10 out of 1500 candidate models, Supplementary Table 1) received sound inputs that were already filtered by the pinnae, head, and torso (head-related transfer functions, HRTFs), and further filtered by the auditory nerve in cochlea. Since these models exhibit many features of human spatial hearing, analyzing their neural representations can bridge the gap between human sound localization behaviors (usually lack of neural data) and auditory systems (Fig. 4).

In real-world listening, there is always noise and reverberation from the environment (Fig. 4a). Saddler & McDermott (2024) simulated a localization-in-noise experiment in which listeners reported which of nine loudspeakers (2 m away, spanning -80° to 80° azimuth in 20° steps) produced a speech utterance, with threshold-equalizing noise played from the remaining eight loudspeakers (top, Fig. 5a). Although the sound azimuth is the same, reverberations blurred the sound waveforms and spectrograms (bottom, Fig. 5a). Therefore, if the sound localization model only needs to disentangle the sound locations, then it should be invariant to changes in content, regardless of whether it is clear or blurred. However, this is not the case: representations of sounds at the same locations but with different content are disentangled but still connected (dots of the same color with and without black centers, Fig. 5b). Furthermore, the sound azimuths on the left side, right side, and at the midline are also disentangled. Our results are consistent across all ten model architectures (Supplementary Fig. 5a) and are robust to different hyperparameter values of UMAP in two model architectures (Supplementary Fig. 6a, b). In summary, “where” models learn to represent sound locations by connecting sounds with two different contents at the same location and also represent nearby sound azimuths closer to each other. They form well separated manifolds but not a map. The “where” model also distinguishes the sound contents by forming two untangled but connected clusters.

To quantify the distribution of nine sound azimuths, we measured the Euclidean distance from each cluster within the same room condition to its nearest neighboring cluster. Fig. 5b (right) shows that the midline azimuth (cyan dots) has the largest distance (13, arbitrary units in UMAP) to its closest neighbor at 20° azimuth (blue dots). The minimal distance (7) to 20° azimuth is at 40° azimuth (violet dots). Across ten model architectures and four UMAP hyperparameter settings, we consistently observed that minimal cluster distances were largest at the midline and its adjacent azimuths (Fig. 5c, left; Supplementary Fig. 5b). Furthermore, this V-shaped dependence of cluster distance on source azimuth was specific to the natural reverberant condition. In the anechoic condition, the curve flattened beyond ±20° azimuth. These geometric patterns observed in the reverberant condition were robustly preserved in the original 512D space using both Euclidean (Supplementary Fig. 5c) and cosine distances (Fig. 5c, right). In contrast, 512D Euclidean distances in the anechoic condition exhibited a flat profile with notably higher variability across models. These results suggest that the object manifolds formed under natural reverberation possess a sufficiently low intrinsic dimensionality that their quantitative structure is preserved across dimensionality reduction (2D vs. 512D) and metric choice (Euclidean vs. cosine).

Importantly, the V-shaped curve of cluster distance is aligned with human behavior, where discrimination of sound azimuth is most sensitive at the midline (largest d’, Fig. 5d, left) and gradually decreases at more lateral azimuths. In contrast, the shape of the model’s task performance curve is flat at the bottom, i.e., it is insensitive to azimuth changes from the midline out to about ±50° (Fig. 5d, right). Thus, the object manifolds provide a better alignment with human behavior than task performance. More importantly, this midline-sensitive representation also supports two-channel models of sound localization, in which azimuth is decoded by the slope of tuning curves rather than by their peak value (McAlpine et al., 2001; Grothe et al., 2010; Chen & Song, 2024) (see Discussion). Together, “what” and “where” manifolds in the models not only align better with behavior than model accuracy, but also help us understand why the models are aligned with human perception.

Saddler & McDermott (2024) showed that precise temporal coding is necessary for sound localization in the horizontal plane. Although we mainly used models with preserved temporal precision (IHC 3000: inner hair cells with a 3000 Hz low-pass cut-off frequency), we also tested ten models with a 50 Hz cut-off frequency (IHC 50). Models with lower temporal precision exhibit lower localization accuracy in the reverberant room even with high SNRs (Supplementary Fig. 5d), but the mechanism is unclear. Our analysis shows that in reverberant conditions, manifolds of sound locations that are far away from the midline overlap (arrows, Supplementary Fig. 5e). This explains why localization accuracy drops even with high SNRs. This is consistent with human behavior, as human localization is most accurate near the midline (Fig. 5d). Furthermore, the representations of locations away from the midline are separated instead of connected between anechoic and reverberant conditions (arrowheads, Supplementary Fig. 5e). Other representation properties are consistent with the high temporal resolution condition. Together, reverberation and lower temporal precision affect the separability of object manifolds of sounds away from the midline.

Our previous analysis shows that, in the horizontal plane, adding multiple background sound sources in reverberant rooms makes their “what” manifolds separate from, but still connected to, those of single foreground sounds in the anechoic room. Thus, the sound localization models are sensitive to changes in spectral content even when these changes are not required for discriminating sound azimuths (which depend on binaural cues, Fig. 4b, left). Most animals, including humans, rely on spectral cues to discriminate sound elevations (peaks and troughs in the HRTFs, Fig. 4b, right). In humans, however, perception depends on relatively coarse spectral features—the transfer function can be smoothed substantially before human listeners notice abnormalities, for reasons that are unclear (Kulkarni & Colburn (1998), Fig. 6a, b). To test whether the trained networks exhibited a similar effect, Francl & McDermott (2022) presented sounds to the models with similarly smoothed transfer functions and measured the extent to which localization accuracy was affected. The effect of spectral smoothing on the networks’ accuracy was similar to the measured sensitivity of human listeners (Fig. 6c), as both showed significant changes at the elbow point (blue squares at 64 cosines). However, the underlying neural mechanisms, in both humans and models, are still unknown.

In order to discriminate sound elevations, the sound localization models need to be sensitive to spectral changes. Our previous analyses of gerbil vocalizations (Fig. 3a) and reverberation (Fig. 5b) in the horizontal plane clearly showed that the “where” model is very sensitive to “what”, regardless of whether the inputs are audio waveforms or cochleagrams. Therefore, we hypothesize that smoothing the original HRTFs with eight different numbers of cosines (from 128 to 1, Fig. 6a) would generate eight different groups of “where” manifolds. Surprisingly, we found that the object manifolds matched human behavior and model performance, but not the sound contents. Compared with the original 256 cosines (white circles), this “what” manifolds of smoothed HRTFs at 64 cosines (silver squares) are fully overlapped with the original, regardless of model architecture (Fig. 6d).

We observed that the “what” manifolds begin to diverge from the baseline manifolds when the number of cosines is smaller than 64. Figure 6d shows the cluster distance between the original HRTFs and conditions with smoothed HRTFs (16 cosines, gray diamonds; 1 cosine, black pentagons) across seven elevations. We quantified these shifts using Euclidean distance in the projected 2D space (Fig. 6e, left) as well as both Euclidean and cosine distances in the native 512D space (Supplementary Fig. 6c; Fig. 6e, right). All metrics consistently demonstrated that manifold separation becomes prominent around 64 cosines, with distances increasing monotonically as spectral density decreases. A notable exception occurred in the 2D Euclidean analysis, where the distance saturated after 4 cosines. Due to the lack of human behavioral data beyond 8 cosines, it remains unclear whether this saturation reflects a projection artifact. Nevertheless, the broad consistency between the 2D and 512D analyses further confirms the low intrinsic dimensionality of these object manifolds.

Collectively, the “where” model exhibits expected sensitivity to large spectral deviations in elevation representations, reinforcing the conclusion that “what” (spectral content) and “where” (location) are inextricably linked. Crucially, the model also mirrors human listeners in its unexpected insensitivity to small spectral changes. This alignment suggests that such robustness arises not merely from acoustic properties, but from the computational nature of the perception task itself. Our analysis thus provides a candidate neural mechanism explaining this perceptual insensitivity.

The sound content used in our previous analysis consists of complex sounds like animal vocalizations, human speech, and noise filtered by the HRTFs. Although they resemble the real-world sounds heard by animals or humans, they preclude us from explaining our findings. Next, we turned our attention to simple sounds that vary in bandwidth and cut-off frequency in the horizontal (Fig. 7) and vertical (Fig. 8) planes. Saddler & McDermott (2024) compared their task-optimized models with human listeners, which allows us to compare our object manifolds to human listeners. In the horizontal plane, human listeners make fewer localization errors when the bandwidth of noise bursts increases (Fig. 7a, b). As shown in Fig. 4b, human listeners mainly rely on binaural ITD and ILD cues for horizontal sound localization. ILDs are significantly affected by the geometry of the head, outer ears, and shoulders, which is why the patterns of the ILD are much more irregular than the ITD patterns (Schnupp et al., 2011). ITDs only exist at low frequencies (Brughera et al., 2013) and are nearly spherically symmetric around the interaural axis (Fig. 7c). Therefore, we hypothesize that low-frequency sounds that preserve ITD cues may help the “where” model form an auditory space map. However, their bandwidth should be neither too narrow (failing to learn structured representations due to low task performance) nor too broad (producing highly untangled manifolds for better task performance). Surprisingly, the “what” and “where” manifolds confirmed our two hypotheses. Broader bandwidth noise bursts with higher task performance form well-separated manifolds (dots with black centers, Fig. 7d). In contrast, the manifolds of most, but not all, narrow-band noise bursts form organized space maps. Why only some of them but not all? We found that this is due to the center frequency: high-frequency sounds (over 1.4 kHz, Brughera et al. (2013)) with unmeasurable high ITD thresholds form clusters, whereas manifolds of low-frequency sounds with low ITD thresholds form maps (Fig. 7e). Our results are consistent across models (Supplementary Fig. 7).

**Figure 7:**
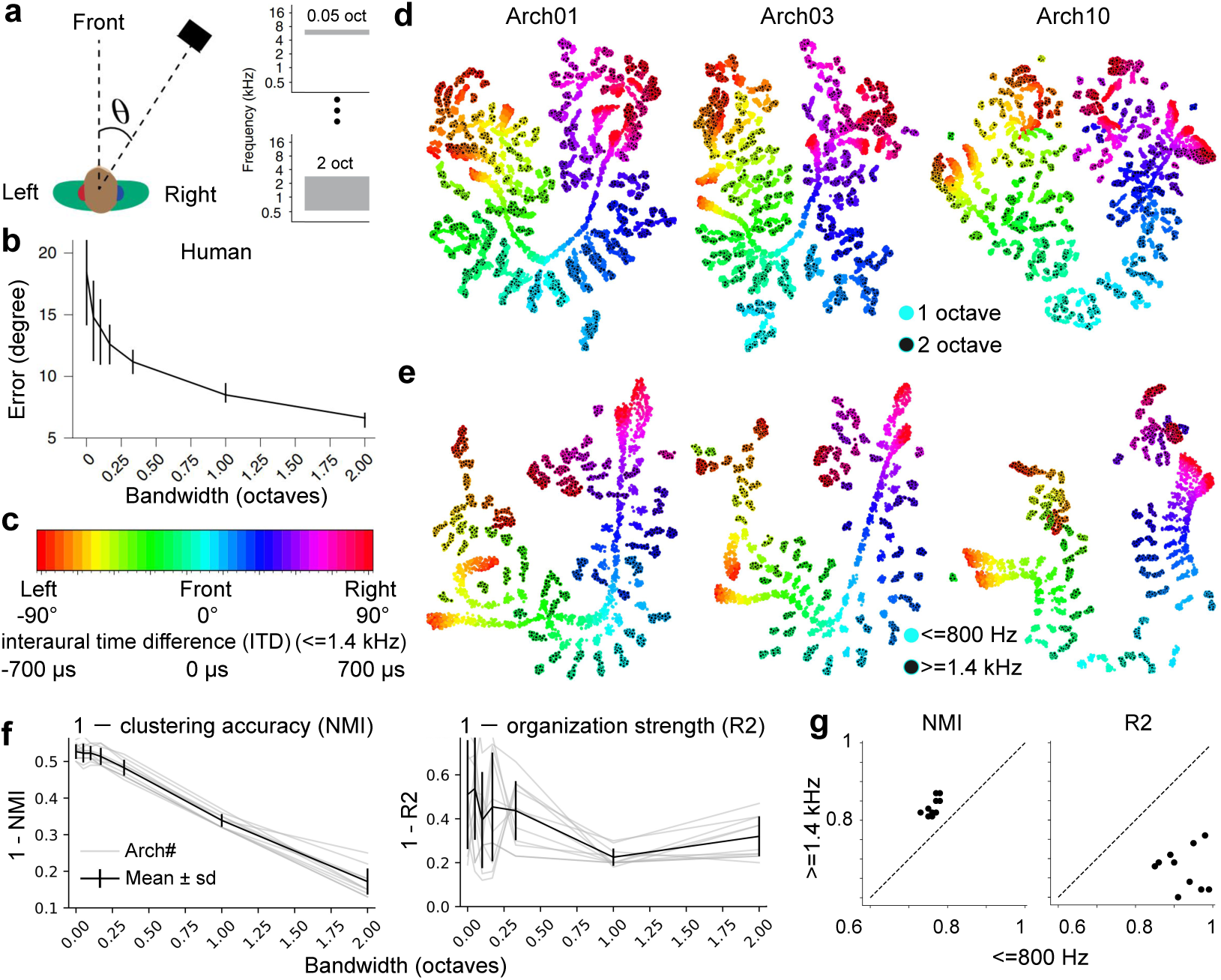
Object manifolds of azimuth are bandwidth- and frequency-dependent and aligned with human behaviors. **a, b**. Schematic of stimuli from an experiment measuring the effects of bandwidth and frequency on localization accuracy. Noise bursts varying in bandwidth and frequency are presented at particular azimuth. Human listeners report the azimuthal position with a key press. Error bars indicate the standard deviation across multiple listeners (copied from Fig. 3d of Francl & McDermott (2022)). **c**. HSV color bar indicates sound azimuth from left to right and ITDs from ipsilateral to contralateral. **d, e**. Representations of 37 sound azimuths at two bandwidths and two cut-off frequencies (bandwidth: 1 octave). **f**. The clustering accuracy and organization strength as a function of bandwidth. We used 1-NMI and 1-*R*^2^ in order to match the human errors in **b**. Thin gray lines: individual model architecture. Thick black line: mean ± one standard deviation. **g**. Scatter plot of NMI and *R*^2^ at two cut-off frequencies. Each dot indicates one model architecture.

**Figure 8:**
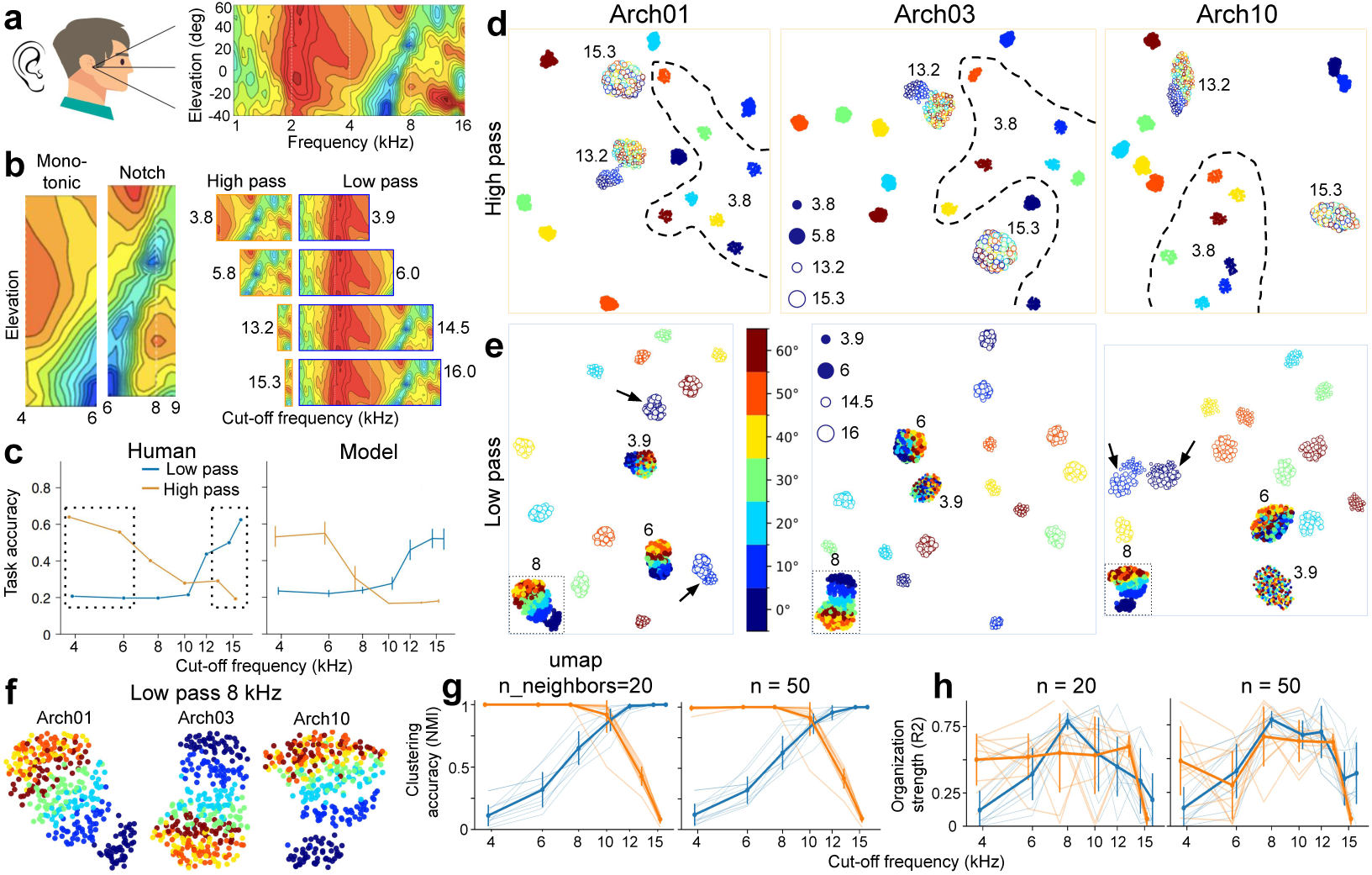
Object manifolds of elevation are spectral-cue dependent and aligned with human behaviors. **a**. Spectral cues available to human listeners at different elevations (copied from Fig. 1 of Zonooz et al. (2019)). **b**. Left: two regions in the HRTFs where the spectral cues vary monotonically with elevation. Right: remaining regions in the HRTFs with four different high- and low-pass cut-off frequencies (dashed squares in **c**). **c**. Effect of cut-off frequencies on localization accuracy in humans and models (copied from Fig. 4n, o of Francl & McDermott (2022)). **d**. Representations of seven elevations at four high-pass cut-off frequencies in three model architectures. **e**. Similar to **d** but for the four low-pass cut-off frequencies. Arrows point to overlapping manifolds from two frequencies at the same elevations. **f**. Elevation maps at an 8 kHz low-pass cut-off frequency. **g**. Clustering accuracy at different high- or low-pass cut-off frequencies with two hyperparameter values in UMAP. Error bars indicate standard deviation. **h**. Similar to **g** but for organization strength.

To quantify our findings and compare them with human behaviors, we plotted the changes in (1 -) clustering accuracy and organization strength against different bandwidths (Fig. 7f). We found that, when increasing the bandwidth in the range where human listeners exhibit low localization error, the clustering accuracy NMI also increased accordingly. Interestingly, the organization strength *R*^2^ increased and reached a peak at 1 octave, and then decreased again from 1 to 2 octaves. This reflects the fact that, on one hand, sound representations cannot remain random when task performance is high, and on the other hand, at the highest performance they tend to form clusters rather than a map. Our results are consistent when testing hyperparameter values in UMAP (Supplementary Fig. 8a). To quantify the bandwidth- and frequency-dependent representations, we compared the clustering accuracy and organization strength of all ten model architectures. The NMIs are always higher when using broader bandwidths or higher center frequencies (Fig. 7g; Supplementary Fig. 8b, c).

Together, the datasets with simple sound stimuli show that object manifolds in the “where” model are sensitive to simple attributes of sound identity like frequency and bandwidth. Importantly, the formation of a map depends on the available ITD cues but deteriorates the model’s and humans’ localization accuracy (*R*^2^ is highest at 1 octave, but accuracy is lower than at 2 octaves).

Our previous analysis shows that when horizontal sound location cues (ITDs) are spatially organized, their corresponding “where” manifolds also form organized maps. Is this organization specific only to horizontal sound locations? As can be seen in Fig. 8a, b, the HRTFs also vary systematically and monotonically with elevation in the 4–6 kHz band (monotonic region), but their changes are markedly smaller than in the 6–9 kHz band (notch region). By contrast, the cues above 9 kHz seem to be strong but are more erratic (non-monotonic). Thus, we hypothesize that a space map would emerge in the vertical plane at specific frequencies and bandwidths of spectral cues.

Saddler & McDermott (2024) compared their task-optimized models with human listeners at different cut-off frequencies and found that they exhibit similar trends: wider frequency bandwidths lead to higher task accuracy (Fig. 8c). Here, we first examined the representations under high-pass conditions with the two smallest and two largest cut-off frequencies (Fig. 8d). Since only a very narrow bandwidth remained after high-pass filtering at 15.3 kHz, the representations tended to be random and not well separated. The representations tended to form a map at 13.2 kHz (with 0° and 10° separated from the others). In contrast, when broad bandwidths were well preserved, the representations formed seven distinct manifolds to represent sound elevations. Importantly, although two sound contents have highly overlapping spectral regions (identical above 5.8 kHz) at the same location, their manifolds never overlapped in any of the ten models (Supplementary Fig. 9). With the low-pass condition, the smaller cut-off frequencies resulted in representations that form a random pattern (3.9 kHz) and a roughly organized map (6 kHz), respectively (Fig. 8e). One piece of evidence for this random pattern is the high dimensionality observed across three model architectures: the bias-corrected PR values were 34.3, 38.6, and 28.9, and the dimensions required for 90% explained variance were 92, 100, and 114. At higher frequencies, the representations of seven elevations formed seven well-separated manifolds. The dimensionality was also very low at 16 kHz (bias-corrected PR: 3.64, 4.18, and 2.51; 90% explained variance: 6, 10, and 26). These results confirm that the unstructured scatter observed in the 2D space is not an artifact of the UMAP, but rather a faithful reflection of the intrinsic high dimensionality and entanglement of the signal itself.

Fig. 8f (also the inset of Fig. 8e) shows the representation at low-pass 8 kHz, where clear space maps are observed. The explanation for this map is the preservation of monotonic and notch regions in the HRTFs, where the spectral cues vary monotonically with elevation. If this is the case, then why does further expanding the bandwidth above 8 kHz make the map disappear? There are two reasons. First, the introduction of extra bandwidth conflicts with the existing monotonic changes. For example, below -20° elevation, the values in the HRTFs are negative around 6 kHz but positive around 12 kHz. The second reason is that forming maps, which causes overlap between nearby elevations, would conflict with the high localization accuracy at broad bandwidths (where forming clusters is the best option). Figs. 8g, h show the quantified results of clustering accuracy and organization strength at two values of the UMAP hyperparameters. The clustering accuracy NMI is consistent with the task accuracy in the human listener and model, i.e., it monotonically increases and decreases for low- and high-pass cut-off frequencies, respectively. In contrast, the trend of organization strength *R*^2^ is different from accuracy, suggesting that whether representations form a map or not is unrelated to accuracy in behavior or in the model. In the low-pass condition, the organization strength peaked at 8 kHz (blue lines, Fig. 8h) and was consistent in all models (Supplementary Fig. 10).

Collectively, the datasets on elevation localization demonstrate that the representations and manifolds in the “where” model are highly sensitive to the available sound content. The formation of a topographic map appears contingent on whether spectral cues contain a single frequency region that varies monotonically with elevation. However, explicit map organization does not guarantee optimal performance in both the models and human listeners.

## 3 Discussion

Here, we examined low-dimensional “what” and “where” representations (or object manifolds) in deep neural network models trained for sound localization. Conceptually, our findings integrate long-standing “what/where” debates with modern representation learning. We built two task-optimized models (five- and ten-layer 1D CNNs) that operate directly on raw binaural waveforms instead of spectrograms/cochleagrams. Recent studies modeling “what” pathways have found that using waveforms as model inputs can even achieve better performance in predicting auditory cortex activity than using the cochleagrams (Li et al., 2023; Tuckute et al., 2023). This usage of raw audio waveforms may not fully capture the complexity of human auditory systems. Therefore, we further analyzed the manifolds of pretrained 2D DNN models from Saddler & McDermott (2024) that were optimized over 1500 architectures, with spectrograms as model inputs. These ten models are equipped with human pinnae/head/torso and cochlea (via HRTFs and auditory-nerve front ends) and trained on human speech under simulated real-world listening environments, including reverberation and background noise. They therefore complement our own models, which involve no architecture search, bear no resemblance to the human auditory system, and are trained on gerbil vocalizations. Thus, our findings are robust across model architectures and datasets.

Our findings are robust across analysis metrics. While we acknowledge that UMAP is a non-linear projection that can potential distort global structure, several lines of evidence validate its utility here. First, the dimensionality of “what” and “where” manifolds is naturally low; even the highest observed dimensionality was less than 10% of the full 512D space, whereas unstructured 2D projections reliably corresponded to high intrinsic dimensionality (Fig. 8e). Second, we observed a high consistency between distances measured in the projected 2D space and the original 512D space (Figs. 5c and 6e), suggesting that the projection preserves essential geometric relationships. Finally, quantitative metrics derived from the 2D space aligned closely with ground truth performance in both models and human listeners (see last paragraph). Collectively, this convergence of low intrinsic dimensionality, consistency with high-dimensional metrics, and alignment with ground truth justifies our use of 2D manifolds for both visualization and quantification.

People may argue that it is completely expected for a “where” model to utilize “what” because it needs to extract sound content information (“what”) in order to localize sounds. This argument is incorrect, because the “what” extracted from the “where” models is not just simple binaural cues like ITDs or ILDs for discriminating sound azimuths. Instead, the “what” consists of complex acoustic features, including gerbil vocalizations with different numbers of harmonics (Fig. 3a), and human speech with and without reverberation (Fig. 5b). People could further argue that separating “what” representations at the same locations only facilitates the “where” model’s localization performance. However, our analysis shows that the representations of “what” are clustered and organized by the spectral similarity. In the vocalization dataset, three harmonic types clustered together, two singleband types formed a separate group, and one broadband noise type lay apart from the single-band group (Fig. 3a). In the reverberant conditions, the manifolds of human speech under anechoic and reverberant rooms are connected but never overlapped (Fig. 5b). People can also argue that the “where” model is simply an acoustic feature extractor, and thus all the representations of “what” will be disentangled. This is also not the case, as the “where” model is insensitive to smaller spectral changes but sensitive to larger spectral changes, with the threshold aligned with human perception (Fig. 6b, e). Together, it is not at all trivial or expected that a “where” model would extract complex acoustic “what” features in order to localize sound locations.

The emergence of “what” in “where” streams suggests a way to resolve the apparent conflict between two dual-stream theories of auditory processing: one emphasizing ventral “what” and dorsal “where” streams (Rauschecker & Scott, 2009), and the other focusing on speech processing (Hickok & Poeppel, 2007). In the latter framework, a ventral stream processes speech signals for comprehension, and a dorsal stream (via Wernicke’s area) maps acoustic speech signals onto frontal articulatory networks. Our results are consistent with the idea that a dorsal “where” model can carry both the content and the location of speech. Our results do not imply that a separate “what” pathway is unnecessary in auditory cortex. Instead, “what” attributes of sounds such as pitch (Bendor & Wang, 2005; Norman-Haignere et al., 2013), voice (Belin et al., 2000; Petkov et al., 2008; Zhang et al., 2024a), speech and music (Norman-Haignere et al., 2015) do form clusters, mainly in the rostral auditory cortex of primates. If DNN models are trained to perform purely “what” tasks or combined “what” and “where” tasks (Saddler et al., 2025), the “what” manifolds should be more untangled (Cohen et al., 2020) than in “where”-only task-optimized models.

Our results suggest that a space map is created by spatially organized localization cues. This is further supported by maps of sound localization cues in the subcortical areas (Olsen et al., 1989; Carr & Konishi, 1990; Ito et al., 2020). Such maps of localization cues may explain why the cue-independent auditory cortex lacks a map of auditory space (Higgins et al., 2017; Wood et al., 2019). We find that the formation of a map does not benefit localization accuracy; instead, it tends to worsen accuracy in both models and human listeners (Fig. 7b vs f; Fig. 8c vs h). The auditory cortex may therefore trade the benefits of forming maps (minimizing wire cost, Chklovskii & Koulakov (2004)) against localization accuracy. A clustered or random pattern instead allows auditory cortex to dynamically represent sound locations under different stimulus contexts (Dahmen et al., 2010; Chen et al., 2025b), top-down attention (Lee & Middlebrooks, 2011; Chen et al., 2025a), or reward (Amaro et al., 2021). The representations of midline azimuths (-20°, 0°, 20°) in the neural manifolds are well separated from other azimuths (Fig. 5c), suggesting that the models encode sound location through sensitivity (slope) rather than the maximally activated location of “where” representation. This is consistent with the two-channel hypothesis of sound localization in both IC (McAlpine et al., 2001; Chen & Song, 2024) and auditory cortex (Stecker et al., 2005; Ortiz-Rios et al., 2017). Notice that we cannot rule out the existence of an auditory space map in the cortex. Since vision can instruct and reshape the auditory space map (Knudsen & Brainard, 1991; King, 1993) or spatial tuning (Jay & Sparks, 1984; Groh et al., 2001) in the IC/SC, we may observe an auditory space map either in audiovisual multisensory cortices (Mazzoni et al., 1996; Schlack et al., 2005) or in models trained for audiovisual multisensory localization tasks (Jia et al., 2025).

Our studies show that analyzing object manifolds allows us to bridge the gap between human sound localization behaviors and the auditory systems, including highest sensitivity to midline azimuths (Fig. 5c, d), insensitivity to small but sensitivity to larger changes in spectral resolution (Fig. 6b, e), and bandwidth-dependent localization accuracy for both sound azimuths (Fig. 7b, f) and elevations (Fig. 8c, g). Francl & McDermott (2022) and Saddler & McDermott (2024) evaluated over ten human sound localization behaviors, but none of them have paired brain recordings. More broadly, it is not practical (e.g., ethical issues) or is impossible (e.g., online Amazon Mechanical Turk) to collect brain data for most human psychophysics experiments. This limitation applies to both audition and vision (Rajalingham et al., 2018; Zhang et al., 2018; Peters & Kriegeskorte, 2021; Dobs et al., 2022; Feather et al., 2023; Sun et al., 2025b; Schulze Buschoff et al., 2025). Fortunately, we have extensive brain data from non-human sensory systems, which enables us to correlate object manifolds in task-optimized models with subjects’ brain activities during tasks. For example, sound localization in the auditory cortex has been investigated across different species for over half a century, and these studies serve as the ground truth of neural manifolds for any computational models. More generally, using DNNs to model sensory systems and human perception, and then analyzing their object manifolds, has widespread potential for understanding the links between brain and behavior. Our studies also show that hypothesis-free models that take real-world stimuli as inputs can learn task-irrelevant “what” manifolds that are aligned with perception. This latent knowledge is often overlooked when we focus only on benchmarking task-relevant alignment (Schrimpf et al., 2020).

## Appendix

### A.1 Code and data availability

Processed data (in .npy format) and code (Python and Jupyter Notebook) to reproduce all figures, except for several diagrams and illustration-only figures (e.g., Figs. 1 and 4), are available at: https://github.com/NeuroscienceAI/WhatWhere. All experiments were run on Ubuntu 22.04.3 LTS with an NVIDIA RTX A5000 GPU and an Intel Xeon W-2225 CPU. We used PyTorch for model training and evaluation. Raw and processed data, including animal vocalizations (Peterson et al., 2024) and human speech (Saddler & McDermott, 2024), were generated and open-sourced by the original authors (see details below). The task-optimized, pretrained models for human speech were also provided by Saddler & McDermott (2024).

### A.2 Method

#### A.2.1 Gerbil datasets and models

We used the Speaker Dataset (Speaker-4M-E1, Peterson et al. (2024)) which was generated by repeatedly presenting five characteristic gerbil vocal calls and a white noise stimulus at three volume levels (18 total stimulus classes) through an overhead tweeter speaker. Between every set of presentations, the speaker was manually shifted by two centimeters to trace a grid of roughly 400 points along the cage floor. This procedure yielded a dataset of 70,914 presentations spanning the 18 stimulus classes. Gerbil vocalizations can range in frequency from approximately 0.5–60 kHz, and different vocalizations correspond to different types of social interactions in nature. In this study, a diverse set of commonly used vocal types was selected that vary in frequency range and ethological meaning. Data is available at: vclbenchmark.flatironinstitute.org.

The network consists of 1D convolutional blocks connected in series (Fig. 2d). It takes raw multichannel audio waveforms as input and outputs the mean and covariance of a 2D Gaussian distribution over the environment. We used two model architectures in this study. One is the simple five-layer DNN model shown in Fig. 2d. The other one is the deeper ten-layer DNN model similar to the DNN model used in Peterson et al. (2024). The size of input channels are 2, 128, 128, 128, 128, 256, 256, 256, 256, and 512. The size of output channels are 128, 128, 128, 128, 256, 256, 256, 256,512, and 512. The kernel size is fixed to 33, and the stride is 1, 2, 1, 2, 1, 2, 1, 2, 1, and 2. We extracted the representations of each sound stimuli (i.e., neural representations) from different layers at the best epoch (minimal validation loss).

#### A.2.2 Human datasets and models

Francl & McDermott (2022) conducted an architecture search over 1500 models and selected the top ten with the lowest validation loss. These models classified noisy 1-second auditory scenes according to the azimuth and elevation of a target natural sound. Source location classes spanned 360° in azimuth (5° bin width) and 0–60° in elevation (10° bin width), yielding a total of 504 output classes (72 azimuth by 7 elevation classes). To ensure that the task was well defined, the training scenes always contained a single natural sound rendered at one target location, superimposed on real-world noise textures diffusely localized at 3–12 different distractor locations. Target sounds were drawn from the Glasgow Isolated Sound Events (GISE-51, Yadav & Foster (2021)) subset of the Freesound Dataset 50k (FSD50K, Fonseca et al. (2021)), which consists of variable-length recordings of individual sources spanning 51 categories of everyday sounds.

To spatialize scenes, Saddler & McDermott (2024) used a virtual acoustic room simulator (Shinn- Cunningham et al., 2001) to render sets of binaural room impulse responses (BRIRs) for a KEMAR manikin in 2000 unique listener environments. The simulator used the image-source method and incorporated KEMAR’s HRTFs (Gardner & Martin, 1995). They randomly generated 2000 unique listener environments by sampling different shoebox rooms (varying in size and wall materials) and listener positions (x, y, z coordinates and head angle) within each room. Room lengths and widths were sampled log-uniformly between 3 and 30 m, and room heights were sampled log-uniformly between 2.2 and 10 m. The listener’s head position was sampled uniformly within each room, with the constraints that the head was at least 1.45 m from wall and no higher than 2 m above the floor.

For each listener environment, they rendered BRIRs at 1008 source locations (2 distances from the listener, 72 azimuths, 7 elevations). One of the distances was fixed at 1.4 m for every BRIR, and the other distance was independently sampled for each BRIR (drawn uniformly between 1 m and 0.1 m less than the distance from the listener to the nearest wall). A total of 1800 unique listener environments were included in the training set, and the remaining 200 were used for validation. The final training and validation datasets consisted of 1,814,400 and 201,600 binaural auditory scenes, respectively. Target natural sounds were placed once at each of the 2000 × 1008 source locations to ensure that the dataset was balanced across the 504 target location classes. Auditory scenes were presented to the model during training at sound levels drawn uniformly between 30 and 90 dB SPL.

We downloaded the open-source model weights and sound stimuli (https://github.com/msaddler/phaselocknet torch) from the authors’ provided Google Drive link. We used the pretrained models in PyTorch versions (simplified IHC 3000 Hz). We used IHC 50 Hz models only in Supplementary Fig. 5d, e. Since the pretrained models are only provided in TensorFlow version, we converted them to PyTorch version with their provided code. We used simplified cochlear model because state-of-the-art cochlear models that best capture the response properties of the auditory nerve are computationally expensive (12 TB). Saddler & McDermott (2024) found that greatly simplified cochlear stage qualitatively and in most cases quantitatively replicated the results obtained with the highly detailed model of the auditory nerve.

In each category of models (simplified or not, IHC 50/320/1000/3000 Hz), there are ten model architectures (arch 01 to arch 10, Supplementary Table 1). We showed the results of architecture 01, 03, and 10 in the Figs. 5678 since they represent CNN with medium (8), deep (10), and shallow (4) layers (Supplementary Table 1). We also showed the results of all ten model architectures in the Supplementary Figs. Those ten models are evaluated on ten different sound localization experiments and we choose four representative experiments: anechoic vs reverberation (Fig. 5) in the horizontal direction, spectral resolution (Fig. 6) in the vertical direction, and bandwidth and frequency dependent sound localization in the horizontal (Fig. 7) and vertical (Fig. 8) directions. We used the sound stimuli from “evaluation” datasets (363 GB), and we used four out of ten datasets: speech in noise in reverb (8.6 GB), spectral smoothing (52.5 GB), bandwidth dependency (19.6 GB), and mp spectral cues (6.9 GB).

Figure 5: anechoic vs reverberation. There are 18,800 stimuli, including 376 speech excerpts, 10 SNRs (-24, -18, -12, -6, 0, 6, 12, 18, 24 dB, plus inf), and 5 rooms (“index room”: 0, 1, 2, 3, 4). The index room = 1 was anechoic room without reverberations. The index room = 0 was the reverberant room Saddler & McDermott (2024) featured in the paper. The remaining three reverberant rooms (2, 3, 4) have low, medium, and high levels of reverberation, but did not use authors in the original paper. We used all four reverberant rooms in this study and did not distinguish between them, therefore there are four times of data points in reverberant rooms than anechoic room (Fig. 5b).

Figure 6: spectral resolution. There are 74,340 stimuli, including 180 azimuths (360° surrounds the subject, uneven distributed), 7 elevations (0° to 60° in steps 10°), 9 smooth factors or different number of cosines (1, 2, 4, 8, 16, 32, 64, 128, 256), and sound duration is 2 seconds. We only analyzed the sounds at midline (0° azimuth).

Figure 7: bandwidth and frequency in the horizontal plane There are 55,500 stimuli, including 37 azimuths (-90° to 90° in steps of 5°, 0° elevation), 12 bandwidths (we only analyzed seven of them to match the human experiments: 0, 1/20, 1/10, 1/6, 1/3, 1, 2), and 125 frequencies (either pure tone or center-frequency of noise burst).

Figure 8: Low-pass and high-pass cut-off frequency of spectral cues There are 9,800 stimuli, including 2 azimuths (front and rear midline), 7 elevations (0° to 60° in steps 10°), 700 combinations of different low-pass (3.9, 6.0, 8.0, 10.3, 12.0, 14.5 or 16.0 kHz) and high-pass (3.8, 5.8, 7.5, 10.0, 13.2 or 15.3 kHz) cut-off frequencies.

We extracted the representations of sound stimuli after passing them through each pretrained model. In each simplified version of DNN model, it contains two parts: one is Peripheral Model which contains Gammatone filter bank to process the audio inputs, and it is same among ten model architectures. The other one is Perceptual Model which is different model architectures. We extract the representations (neural representations, 512 dimensions) of each sounds after multiple convolutional layers (depending on model architectures) at the “fc intermediate” (penultimate) layer.

#### A.2.3 Dimensionality estimation and participation ratio

##### Dimensionality Estimation

To quantify the geometric structure of the neural representations, we analyzed the eigenspectrum of the activation patterns. Let **X** ∈ R*^M×N^* denote the matrix of neural activations, where *M* is the number of samples (stimuli) and *N* is the number of neurons (ambient dimension, e.g., 512). We first computed the centered covariance matrix **Σ̂**:

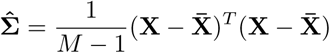

where **X̄** is the mean activation vector across samples. We then obtained the eigenvalues *λ*_1_ ≥ *λ*_2_ ≥ · · · ≥ *λ_N_* of **Σ̂**.

##### Standard Participation Ratio (PR)

We calculated the standard participation ratio (Gao et al., 2017), a continuous measure of the effective dimensionality of the manifold, defined as:

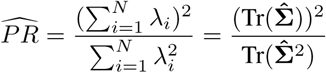

where Tr(·) denotes the trace of the matrix. This metric quantifies how evenly the variance is distributed across dimensions; it approaches 1 if variance is concentrated in a single dimension and approaches *N* if variance is equally distributed.

##### Bias-Corrected Participation Ratio

Recent theoretical work indicates that 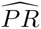 is a biased estimator when the sample size *M* is finite, specifically underestimating the true dimensionality due to the inflation of the second moment term (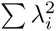) by sampling noise (Chun et al., 2025). To address this, we applied a bias-corrected estimator. We estimated the population second moment by subtracting the contribution of sampling bias:

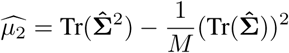

The bias-corrected participation ratio (*PR_bc_*) was then computed using this adjusted second moment:

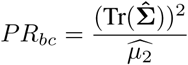

This correction ensures that the estimated dimensionality reflects the underlying manifold structure rather than artifacts of limited sampling.

#### A.2.4 UMAP, NMI, and *R*^2^

We chose UMAP (version: 0.1.1), a nonlinear dimensionality reduction method that is explicitly designed to preserve the geometric structure of high-dimensional data, to reveal object manifolds (McInnes et al., 2018). UMAP has been widely used to uncover neural manifolds in recent years (El-Gaby et al., 2024; Sun et al., 2025a; Zutshi et al., 2025). UMAP preserves both local and global geometry. For local structure, UMAP keeps similar data points close together: if two points are neighbors in the high-dimensional space, they are likely to remain neighbors in the low-dimensional representation. For global structure, distances between distinct clusters in the 2D output are meaningful and reflect the relationships among groups in the original high-dimensional space. Two hyperparameters primarily control the representation geometry. n neighbors determines how UMAP balances local versus global structure: low values emphasize local detail, whereas high values capture the global geometry. min dist controls how tightly points are packed in the output space. In our analyses, we varied only the major hyperparameter, n neighbors, and kept all other hyperparameters at their default values (e.g., min dist = 1, n components = 2, random state = 0).

We used the scikit-learn package (version: 1.6.1) to quantify clustering accuracy and organization strength. Clustering accuracy was measured using normalized mutual information (NMI; normalized mutual info score from “sklearn.metrics”). We used KMeans (from “sklearn.cluster”) with the number of clusters set to the number of classes (i.e., sound types or azimuths) to assign cluster labels. An NMI value of 1 indicates perfect agreement between predicted cluster labels and ground-truth labels. Organization strength was measured using explained variance (*R*^2^; r2 score from “sklearn.metrics”) between predicted and ground-truth labels. We used linear regression (from “sklearn.linear model”) to fit the 2D or 3D representations of each stimulus to its ground-truth label, and then predicted the label for each stimulus.

#### A.2.5 Distance of manifolds

Centroids for each condition (e.g., specific azimuth or elevation) were computed by averaging the activation vectors of all stimuli belonging to that class. We computed pairwise distances between these centroids using three distinct metrics to capture different aspects of the representation.

##### Low-dimensional (2D) projection distance

We computed the Euclidean distance between cluster centroids in this 2D projection:

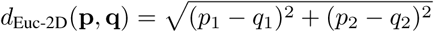

where **p** and **q** are the 2D coordinates of the projected centroids.

##### High-dimensional cosine distance

To measure the representational similarity between conditions independent of overall activity magnitude, we computed the cosine distance in the original 512D space. For two activation vectors **u** and **v**, the cosine distance is defined as:

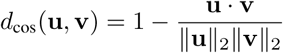

where ||·||_2_ denotes the *L*^2^ norm. This metric relies on the angle between vectors, making it robust to variations in global activations (vector magnitude) and sensitive to the pattern of activation across units. *d*_cos_ ranges from 0 to 2 (0=identical, 1=orthogonal, 2=opposite).

##### High-dimensional Euclidean distance

We also computed the standard Euclidean distance in the 512D space to account for magnitude differences:

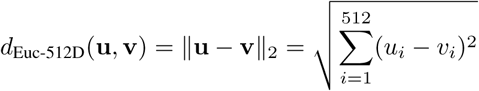

Unlike cosine distance, this metric is sensitive to the total energy of the neural activations.

### A.3 Supplementary Figures and Table

**Supplementary Figure 1:**
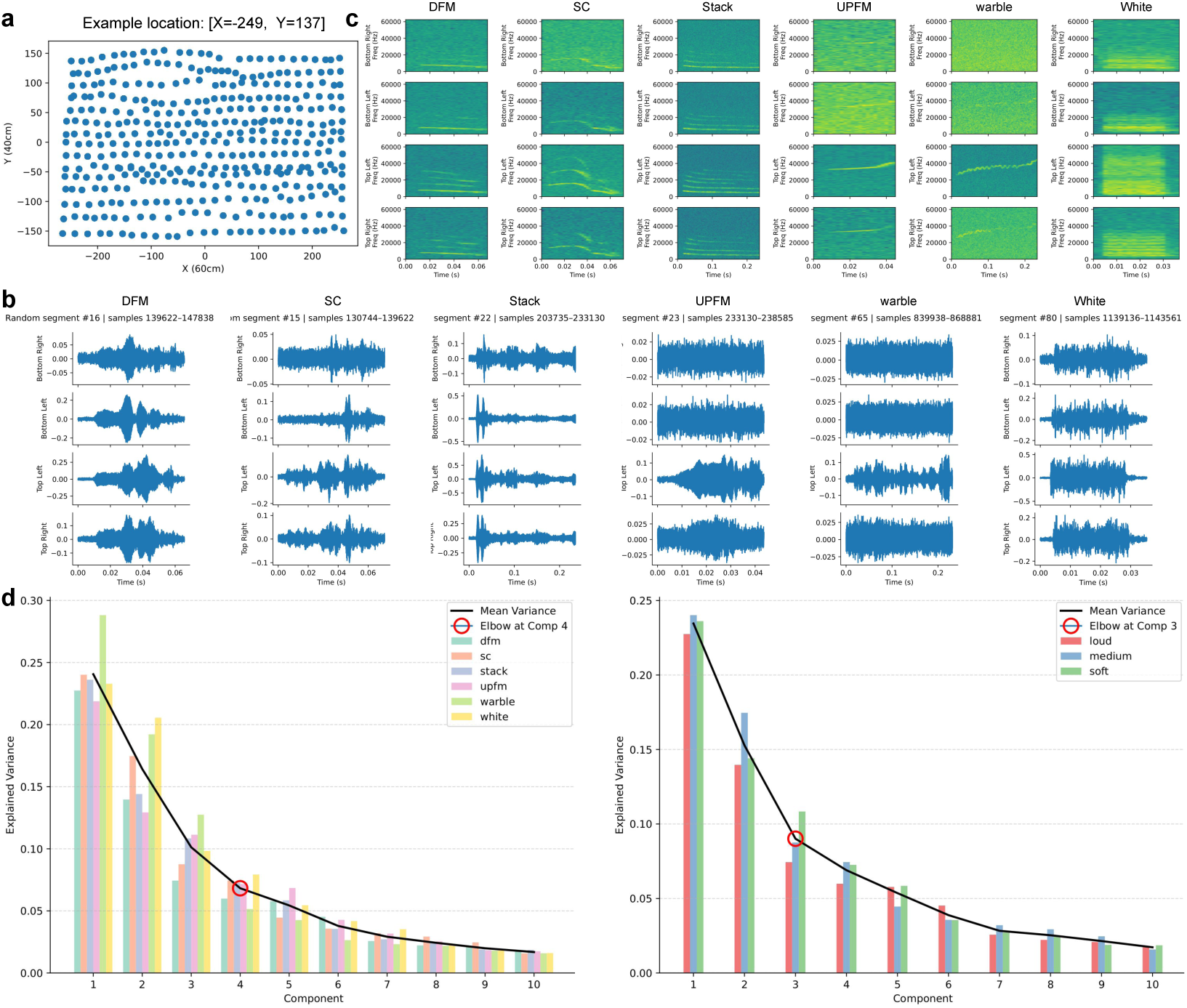
**a**. There are 394 different sound locations, and the example location comes from the top left corner. Each location (big blue dot) has many (i.e., 144) smaller dots inside since different sound types, levels, and a median of eight different samples are presented from there. **b**. Audio waveforms from six sound types (columns) that were recorded by four microphones at four corners. Notice that the amplitudes are different for each plot. **c**. Spectrograms for six types of waveforms shown in **b**. **d**. Explained variance (Y-axis) and number of component/dimension (X-axis) for six sound types (Left) and three sound levels (Right). The averaged *R*^2^ among five sound types across ten different components are 0.24, 0.16, 0.10, 0.07, 0.05, 0.04, 0.03, 0.02, 0.02, 0.02 (black curve). The averaged *R*^2^ among three sound levels across ten different components are 0.23, 0.15, 0.09, 0.07, 0.05, 0.04, 0.03, 0.03, 0.02, 0.02. Red circles highlight the elbow points at 4th and 3rd components.

**Supplementary Figure 2:**
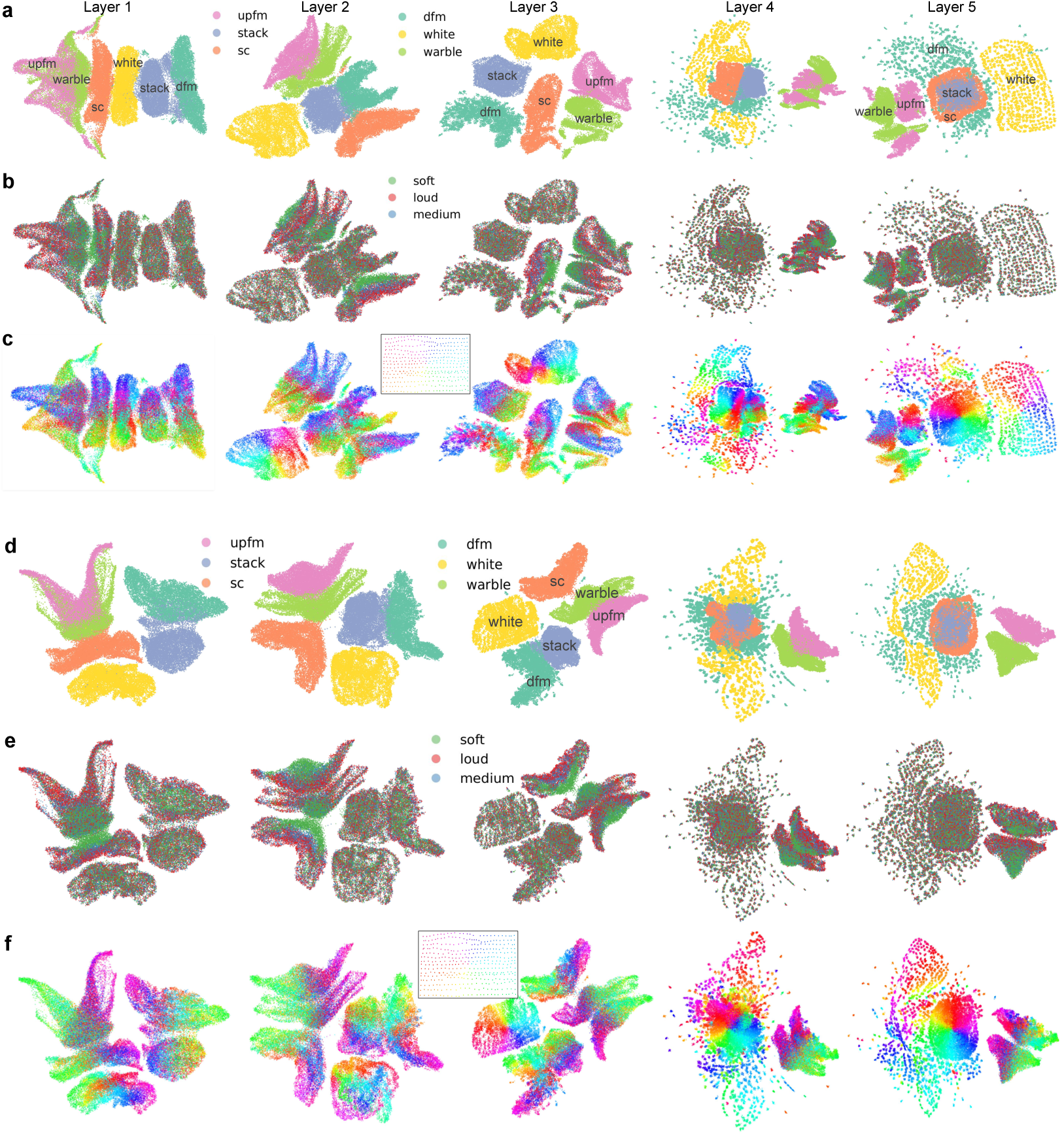
Representations of “what” and “where” attributes of sounds in all five layers. **a-c**. Representations of six sound types, three sound levels, and 394 sound locations. Data in layer 5 was the same as Figure 3a-c. The microphone pair is M24. The NMIs for sound types are 0.5454, 0.6140, 0.8898, 0.5488, and 0.6233. The NMIs for sound levels are 0, 0.0001, 0.0025, 0, and 0. **d-f**. Similar to **a-c**, but for the microphone pair of M1 and M3. The NMIs for sound types are 0.6892, 0.8313, 0.8095, 0.4953, and 0.6582. The NMIs for sound levels are 0.0001, 0.0003, 0.0002, 0, and 0.

**Supplementary Figure 3:**
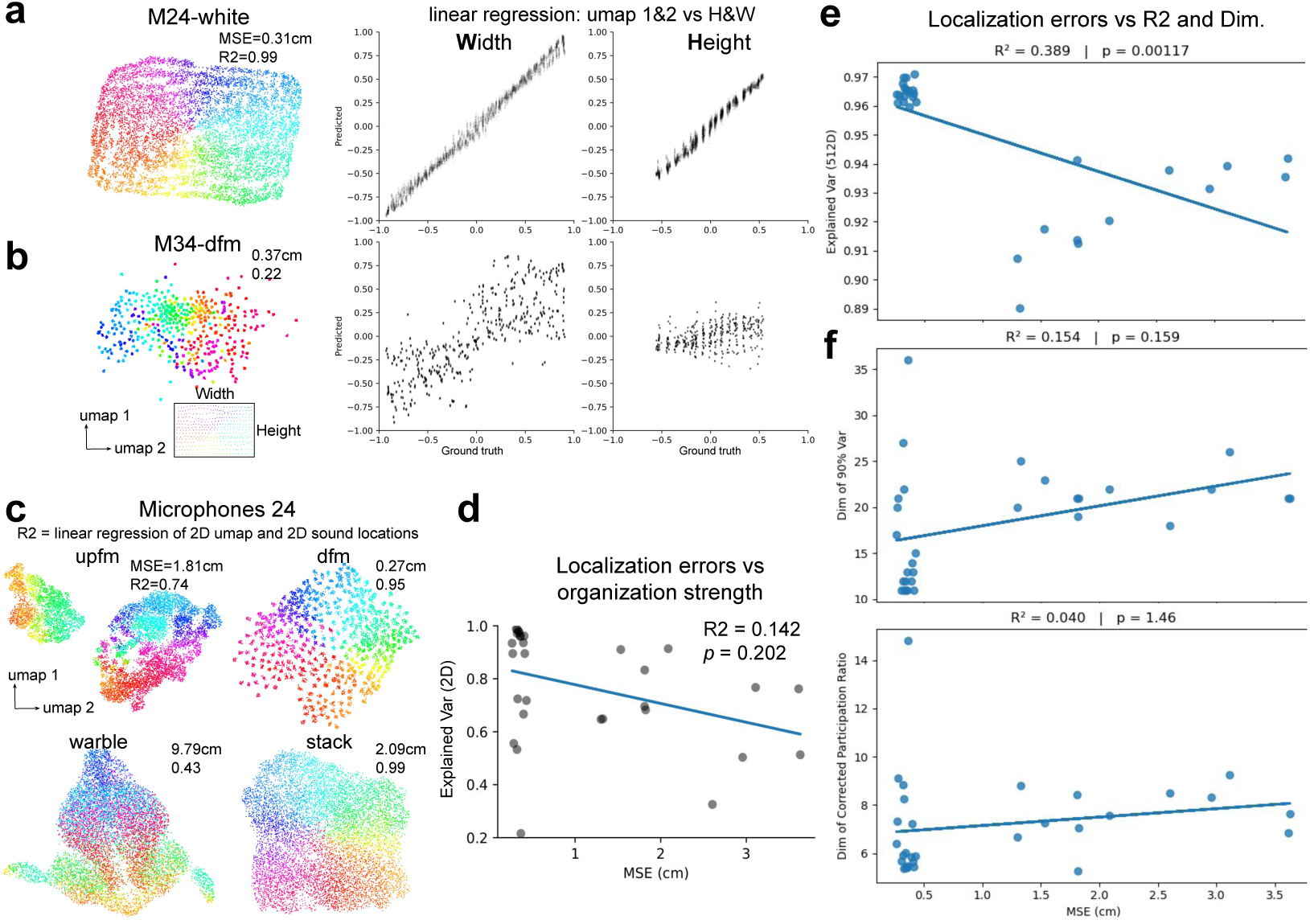
Linear regression of sound location representations and locations of the speaker. **a**. A white noise sound type from the microphone pair of M2 and M4. Left, 2D UMAP visualization of sound location representation. Middle, linear regression of 2D UMAP against the width of speaker location. Right, regression of UMAP against the height of speaker location. **b**. Similar to **a** but for a different microphone pair and sound type. **c**. Same microphone pair as Fig.3c, but sound stimuli from each sound type were trained separately. **d**. Localization errors (MSE) versus explained variance in 2D UMAP. Each dot represents one session (30 total: five sound types × six microphone pairs). Data from the “warble” call type are excluded because their MSE values exceed 10 cm. **e**. Localization errors (X-axis) versus explained variance in raw 512D space. **f**. Localization errors (X-axis) versus number of dimensions needed to explain over 90% variance (Top, Median=19.5, Min=11, Max=36), and effective dimensionality or the participation ratio after bias-correction (Bottom, Mean=7.192, Min=5.272, Max=14.833) (without correction: Mean=7.186, Min=5.269, Max=14.810). The p-values are corrected by multiplying the number of sound types (N=5).

**Supplementary Figure 4:**
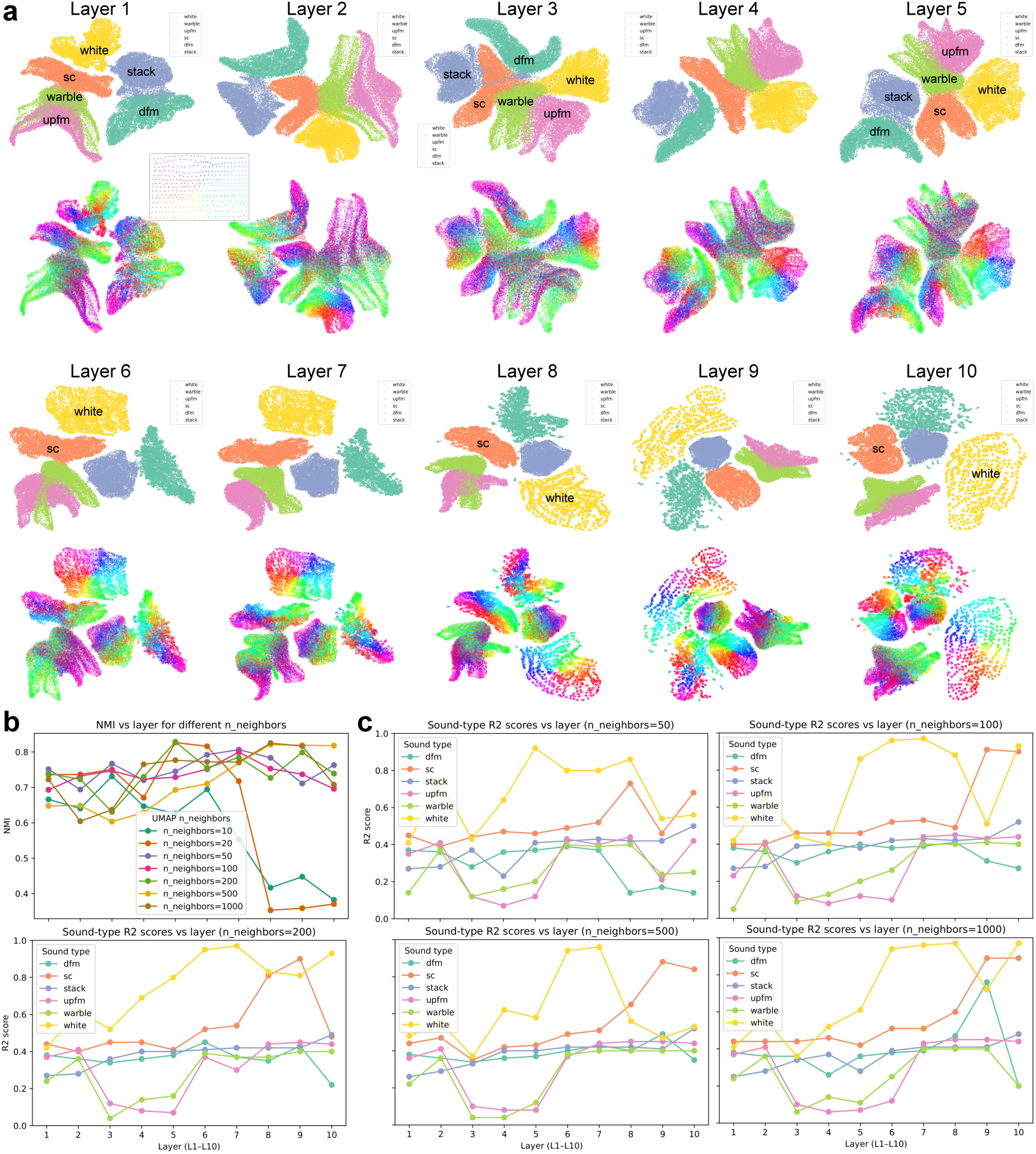
**a**. Representations of sound types (1st and 3rd rows) and locations (2nd and 4th rows) from layer 1 to layer 10 in a deeper CNN for the microphone pair of M1 and M3. The number of neighbors in UMAP is 200. **b**. The NMIs across all ten layers. The microphone pair is M1 and M3. The NMIs panel quantifies the clustering performance when changing the hyperparameter (number of neighbors) of UMAP. **c**. The *R*^2^ scores across all ten layers. Notice that the NMIs drop suddenly after layer 6 when the number of neighbors is 10 and 20. Therefore, we only show the *R*^2^ scores of six sound types across ten layers with the number of neighbors larger than 20.

**Supplementary Figure 5:**
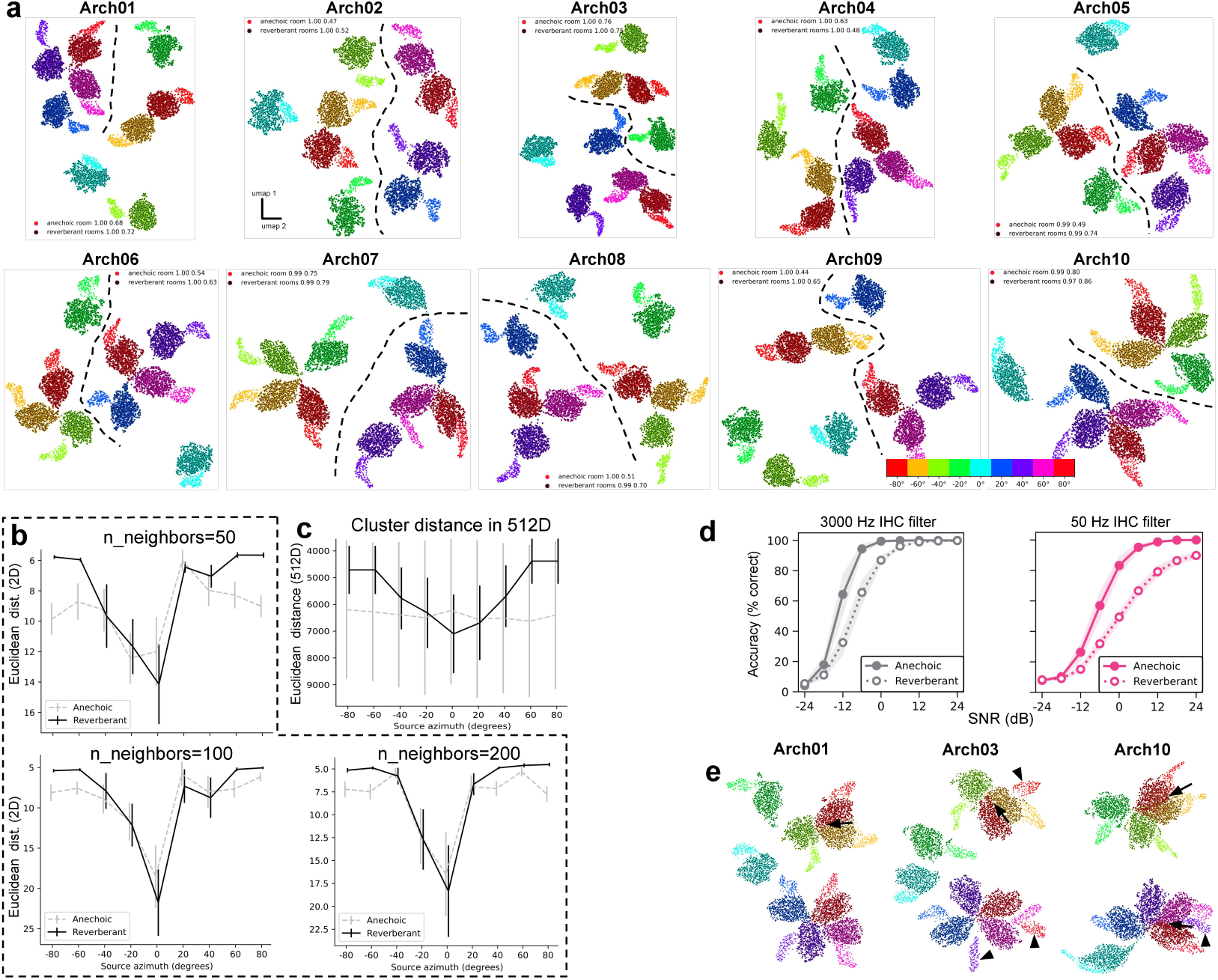
**a**. Representations of nine sound azimuth angles in the anechoic and reverberant rooms in all ten model architectures. The number of neighbors in UMAP is 20. The dashed black lines indicate the manually drawn boundary between four left and four right azimuths. In each figure legend, the first value is NMI (clustering performance), and the second value is *R*^2^-score (organization strength). Three panels (Arch 01, 03, and 10) are shown in Fig. 5b. **b**. The minimal Euclidean distance between manifolds in the 2D space across three different hyperparameter values of UMAP. **c**. The minimal Euclidean distance between manifolds in the raw 512D space. **d**. Model sound localization accuracy as a function of signal-to-noise ratio (SNR) and reverberation. Shaded areas represent the standard deviation across ten model architectures. IHC: inner hair cells. The figures are copied from Fig. 7e in Saddler & McDermott (2024). **e**. Similar to **a** but with a 50 Hz IHC filter. Arrows point to overlapping representations between -80° and -60° or between 60° and 80° in the four reverberant rooms. Arrowheads point to separated representations between anechoic and reverberant rooms at the same azimuths.

**Supplementary Figure 6:**
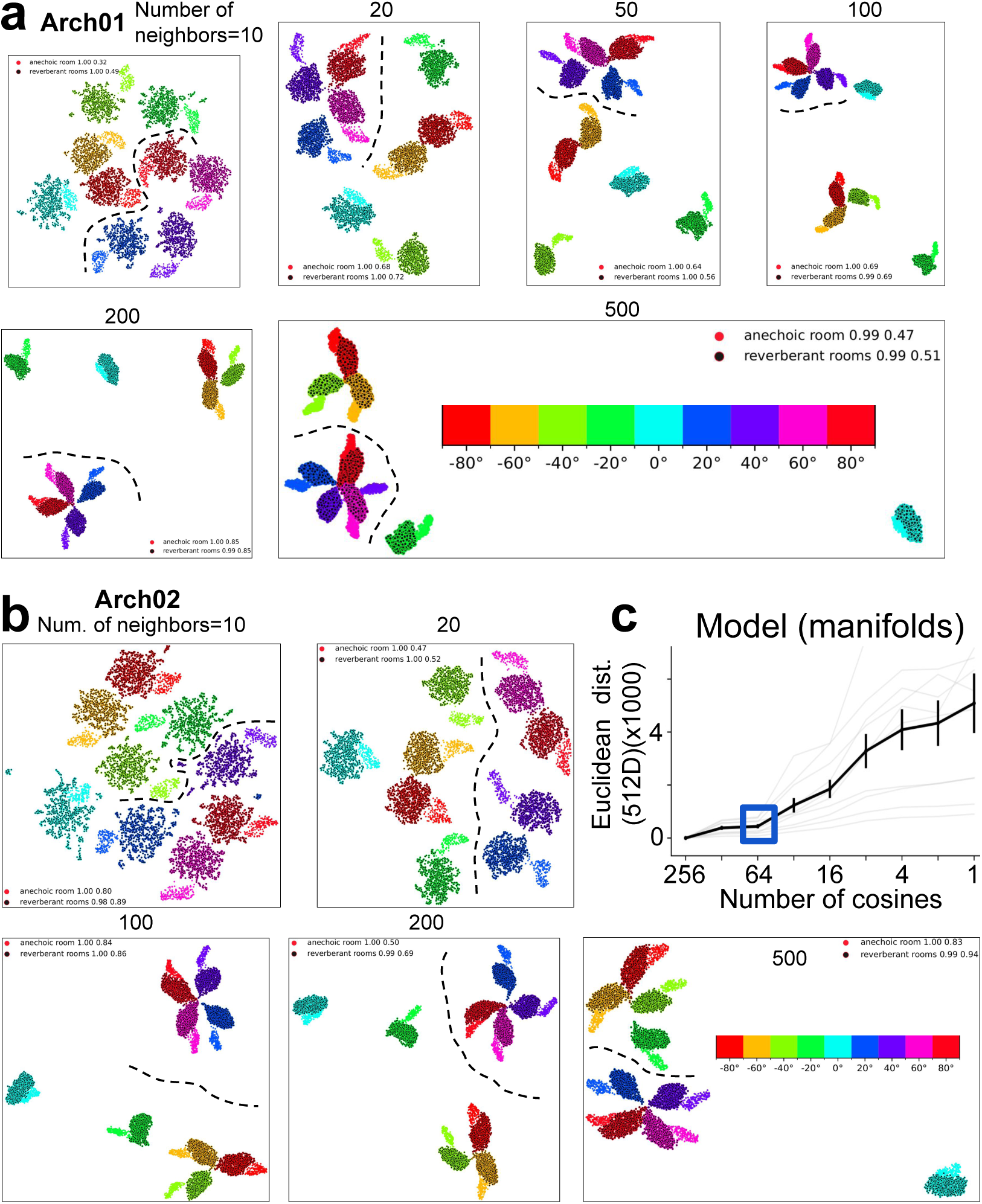
**a**. Representations of nine sound azimuth angles in the anechoic and reverberant rooms with six different hyperparameter (number of neighbors) values of UMAP. The dashed black lines indicate the manually drawn boundary between four left and four right azimuths. In each figure legend, the first value is NMI (clustering performance), and the second value is *R*^2^- score (organization strength). One panel (number of neighbors is 20) is shown in Fig. 5b. **b**. Similar to **a** but with model architecture 02. **c**. The euclidean distance to original 256 cosines (Y-axis) at different number of cosines (X-axis) in the raw 512D space. Blue square highlights the elbow point. This figure is relevant to Fig. 6e.

**Supplementary Figure 7:**
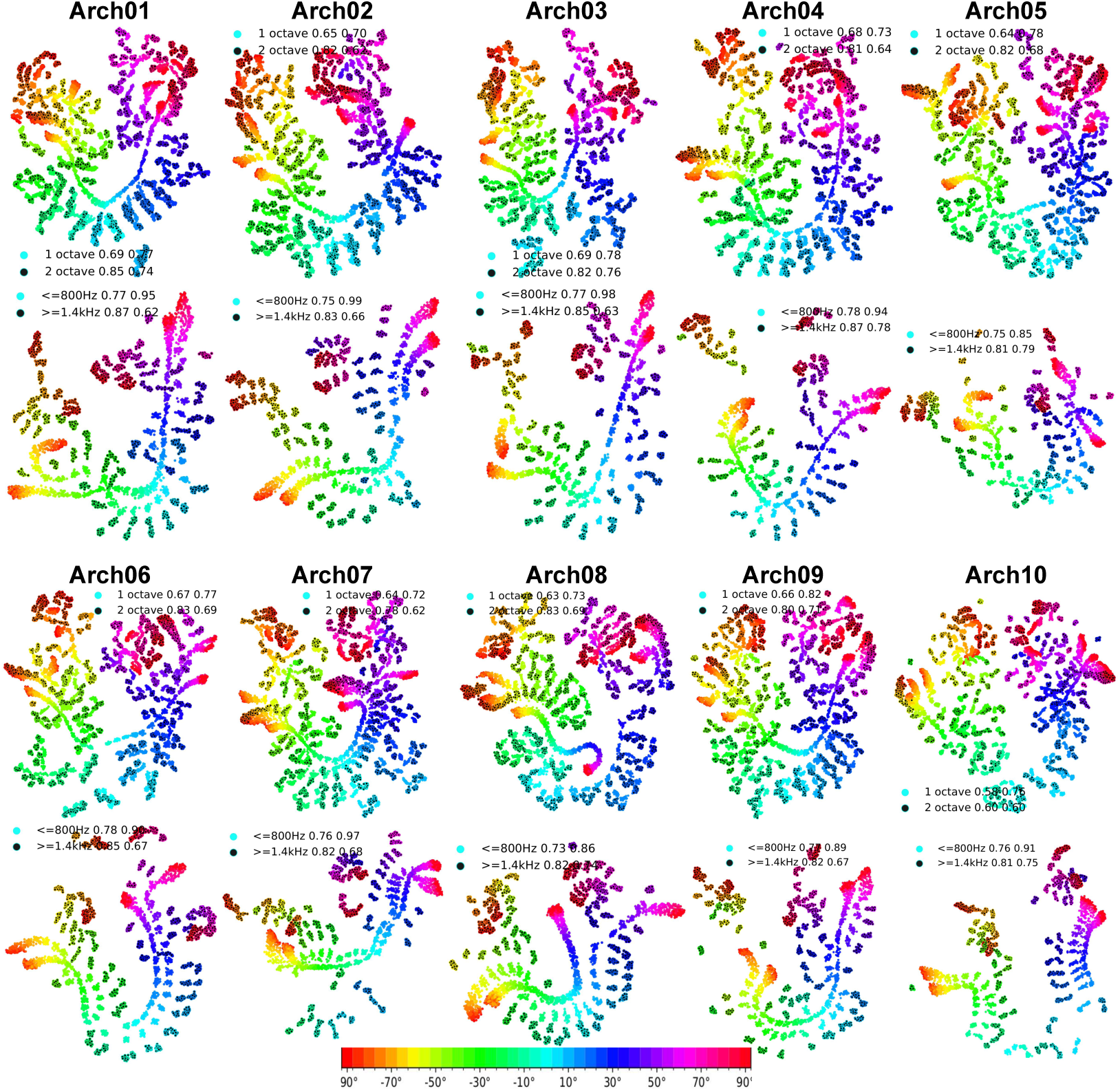
Representations of sound azimuths at different bandwidths (1st and 3rd rows), center frequencies (2nd and 4th rows), and network architectures. Representations in Arch 01, 03, and 10 are shown in Fig. 7d, e. In each figure legend, the first value is either bandwidth or cut-off frequency, the second value is NMI (clustering performance), and the third value is *R*^2^-score (organization strength).

**Supplementary Figure 8:**
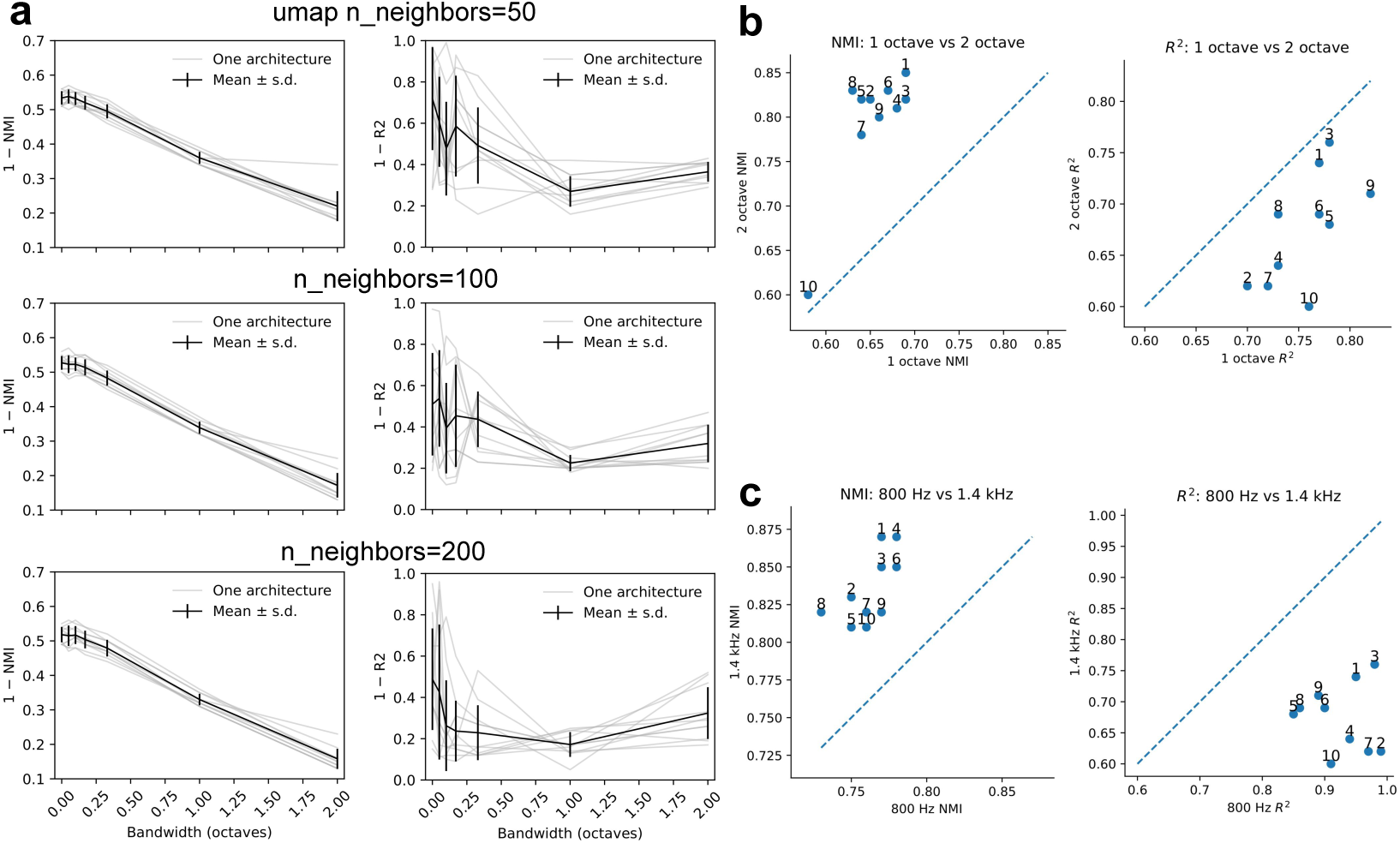
The clustering performance and organization strength at different bandwidth, network architectures, and hyperparameter of UMAP. **a**. Similar to Fig. 7f but with three different number of neighbors of UMAP (first to third row). The second row is same to Fig. 7f. **b**. The scatter plot of NMIs (left) and *R*^2^ scores (right) across ten model architectures. X-axis is 1 octave bandwidth, and Y-axis is 2 octave bandwidth. The value above each blue dot represents each model architecture. **c**. Same data as Fig. 7g. The scale of X and Y axis for the NMIs panel is different from the *R*^2^ scores panel.

**Supplementary Figure 9:**
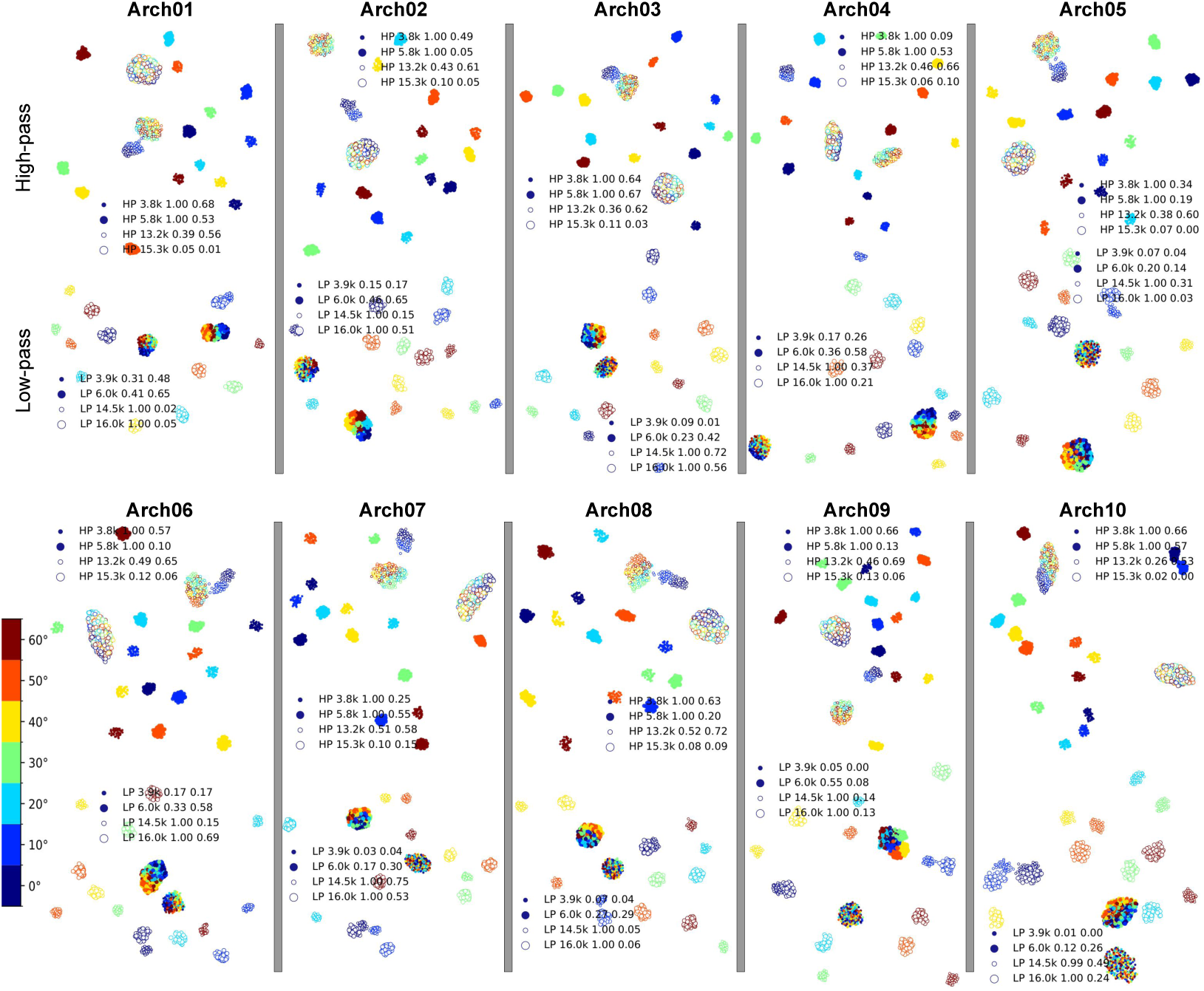
Representations of seven sound elevation angles in all ten network architectures at four different high-pass (HP, 1st and 3rd rows) and low-pass (LP, 2nd and 4th rows) cut-off frequencies. Representations in Arch 01, 03, and 10 are shown in Fig. 8d, e. In each figure legend, the first value is cut-off frequency, the second value is NMI (clustering performance), and the third value is *R*^2^-score (organization strength).

**Supplementary Figure 10:**
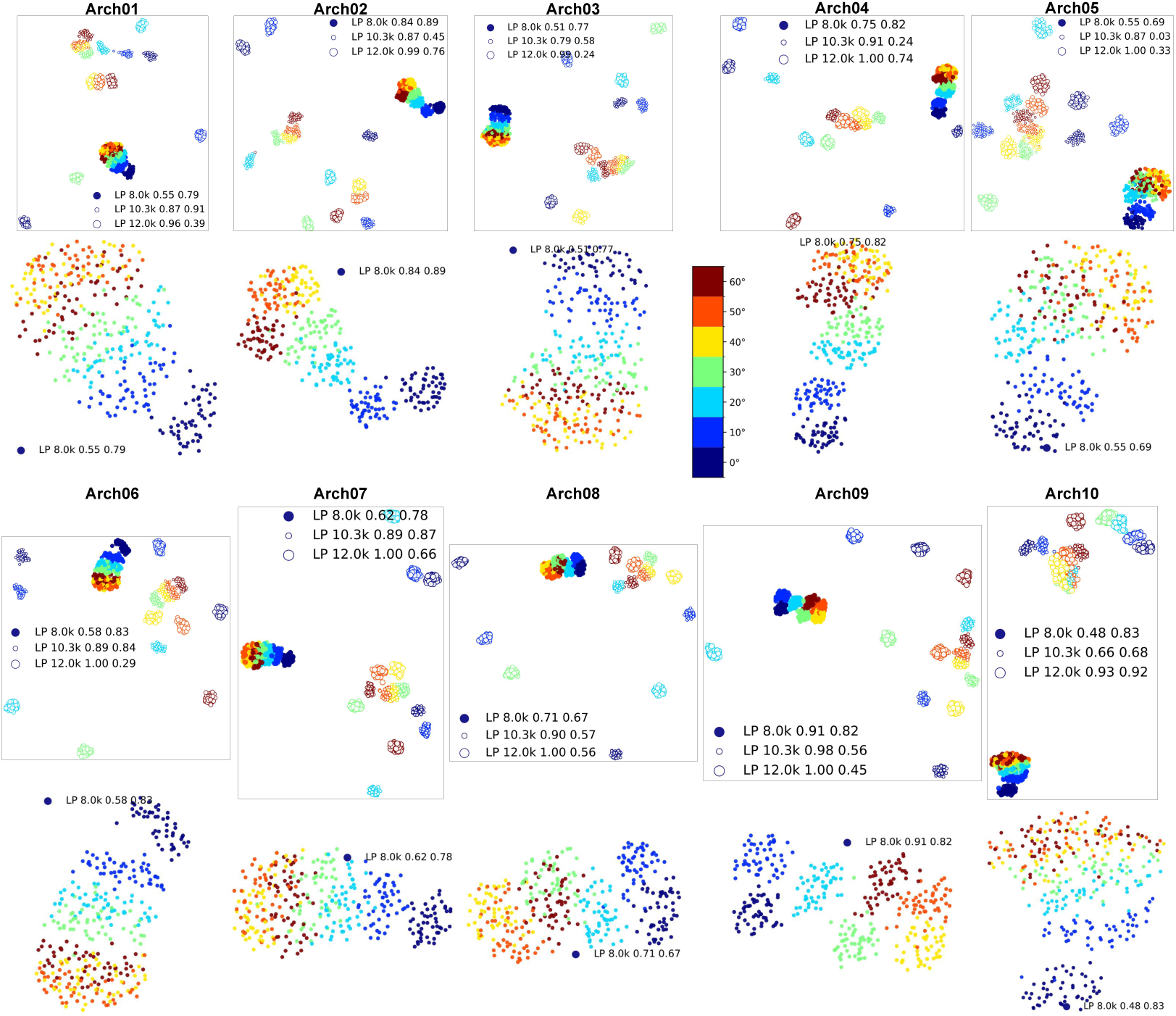
Representations of seven sound elevation angles in all ten network architectures at three intermediate low-pass (LP, 1st and 3rd rows) cut-off frequencies. The 2nd and 4th rows show the same representations but with only one 8 kHz low-pass frequency. Representations in Arch 01, 03, and 10 are shown in Fig. 8f. In each figure legend, the first value is low-pass cut-off frequency, the second value is NMI (clustering performance), and the third value is *R*^2^-score (organization strength).

**Supplementary Table 1.**
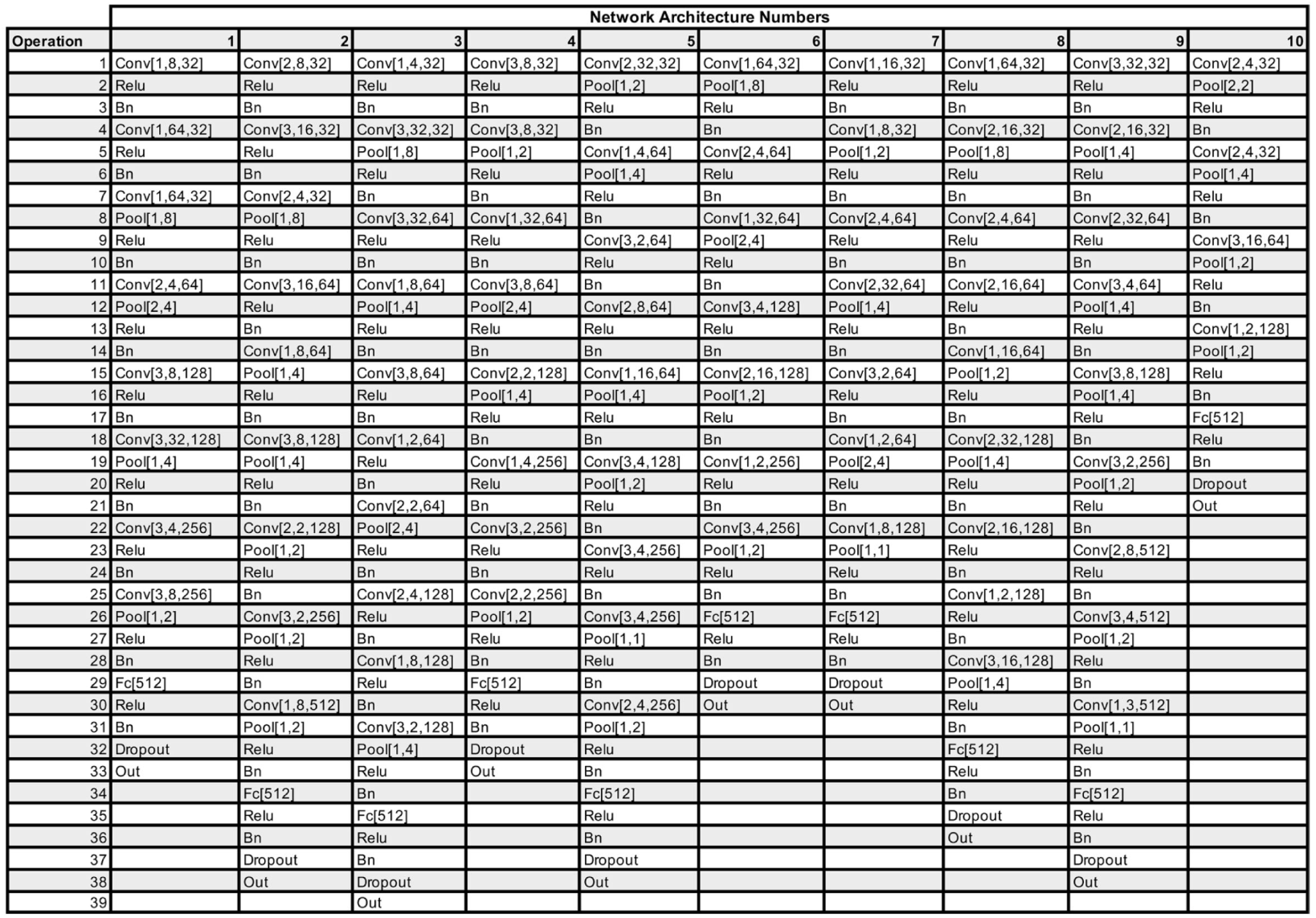
Summary of the 10 network architectures. Conv[X, Y, Z]: Convolutional layer with kernel height X, kernel width Y, and number of filters Z. ReLu: Rectified linear unit layer. Bn: Batch normalization layer. Pool[X, Y]: Max pooling layer with kernel height X and kernel width Y. Fc[X]: Fully connected layer with X number of units. Dropout: Dropout layer. Out: Softmax classification layer with 504 output units. This table is copied from the Extended Data Fig. 3 of Francl & McDermott (2022) (same as Supplementary Table 1 of Saddler & McDermott (2024)).

